# PEACOC - Detecting and classifying a wide range of epileptiform activity patterns in rodents

**DOI:** 10.1101/2024.03.14.584961

**Authors:** Katharina Heining, Antje Kilias, Philipp Janz, Carola A Haas, Ulrich Egert

## Abstract

Epileptiform activity (EA) manifests in diverse patterns of hypersynchronous network activity. Fundamental research mostly addresses two extreme patterns, individual epileptiform spikes and seizures. We developed PEACOC to detect and classify a wide range of EA patterns in local field potentials. PEACOC delimits EA patterns as bursts of epileptiform spikes, and classifies these bursts according to spike load. In EA from kainate-injected mice, burst patterns displayed a continuum of spike loads. With PEACOC, we partitioned this continuum into bursts of high, medium and low spike load. High-load bursts resembled electrographic seizures. The EA patterns automatically retrieved by PEACOC were reproducible and comparable across animals and laboratories. PEACOC has been employed to diagnose the overall burden of EA in individual mice, and to describe epileptic dynamics at multiple time-scales. We here further report that the rate of high-load bursts was anti-correlated to granule cell dispersion in the dentate gyrus.

## Introduction

The two most well studied patterns of epileptiform activity (EA) are spikes and seizures, yet there are other patterns of EA that have received less attention thus far. A tool that can comprehensively quantify the spectrum of such patterns could greatly further our understanding of epilepsy.

Particularly rich patterns of EA are found in mesial temporal lobe epilepsy (MTLE), which is arguably the most common form of adult epilepsy (Engel, 2001; Téllez-Zenteno and Hernández-Ronquillo, 2011). MTLE patients suffer from frequent, non-convulsive seizures, which are often refractory to medication (Chabardès et al., 2005; Engel, 2001). A clear border between the focal seizures and other, subclinical, patterns of epileptiform activity does not exist, and it has been argued that the whole spectrum of EA, ranging from single epileptiform spikes to seizures, is continuous (Gotman, 2011; Sperling and O’Connor, 1990; Walker and Kovac, 2015; Zangaladze et al., 2008a). MTLE is typically accompanied by hippocampal sclerosis, characterized by network restructuring, cell loss and granule cell dispersion (GCD, Blümcke et al., 2009; Walker, 2015). The intrahippocampal kainate mouse model replicates key electrophysiological and histopathological features of MTLE, and is widely used in experimental research (Heinrich et al., 2011; Janz et al., 2017; Krook-Magnuson et al., 2013; Riban et al., 2002; Suzuki et al., 1995). Thus far, EA in this model has been described as consisting of solitary spikes, spike trains, and electrographic seizures, while generalized seizures are very rare (Maroso et al., 2011a; Riban et al., 2002; Twele et al., 2016a).

Hitherto, electrographic seizures in this epilepsy model were distinguished from other patterns of EA mostly by several heuristic criteria and thresholds (reviewed in Twele et al., 2016). Typically, threshold values are set by experts manually annotating the data. This makes detecting electrographic seizures a laborious procedure, the results of which can be difficult to compare across individual animals and experts.

Electrographic seizures can be considered as bursts of epileptiform spikes. Thus, automatically detecting their constituent spikes would solve some of the issues associated with manual annotation. However, decreasing amplitudes and changing waveforms of spikes within seizure-like bursts impede their automated detection based on waveform templates and ampltiude thresholds. Existing spike detection algorithms therefore commonly focus on solitary spikes with high spike amplitudes or clearly discernible waveforms (Anjum et al., 2018; Chauvière et al., 2012; Huneau et al., 2013; Karoly et al., 2016; Krook-Magnuson et al., 2013). Seizure detectors, on the other hand, are mostly designed to specifically detect large, generalized seizures (Esteller et al., 2001; Gardner et al., 2006; Krook-Magnuson et al., 2013; Logesparan et al., 2013). These seizure detectors are often based on measures of signal energy, which can vary substantially depending on signal quality. They therefore generally require subject-specific tuning in order to obtain comparable detections (e. g. Armstrong et al., 2013). The uncertainty associated with the definition of seizures poses a further problem (Gotman, 2011; Walker and Kovac, 2015). Critically, due to the implicit dichotomy between spike detectors and seizure detectors, the wide range of patterns between the extremes of “spike” and “seizure”, which constitutes a large fraction of EA, can easily be overlooked.

To improve EA quantification and classification, we developed an algorithm to comprehensively and automatically detect and classify EA in local field potentials (LFPs). The PEACOC (**P**atterns of **E**pileptiform **A**ctivity: a **CO**ntinuum-based **C**lassification) software detects solitary spikes as well as spikes within dense, seizure-like activity. This allows investigating and classifying a wide range of EA patterns within a single, unified framework. Our results support the hypothesis that EA forms a continuum. PEACOC yields reproducible and robust results, and can be tailored to a variety of purposes. To date, we used it in several studies to establish a connection between EA and anatomical and functional markers of MTLE in the intrahippocampal kainate mouse model (Heining et al., 2019; Janz et al., 2018, 2017; Kilias et al., 2018; E. Paschen et al., 2020; Tulke et al., 2019). Here we provide a detailed description, validation, and explanation of the methodology.

## Results

LFPs were recorded using intrahippocampal electrodes close to the kainate injection site. Freely-behaving mice displayed EA typical of this mouse model (Figure 1A; Riban et al., 2002; Twele et al., 2016). While epileptiform spikes could occur in isolation, most were part of compound events. These varied in size and morphology, ranging from small groups of a few loosely spaced spikes to larger bursts and ultimately events with morphologies indicative of behavioral seizures.

**Figure 1:**
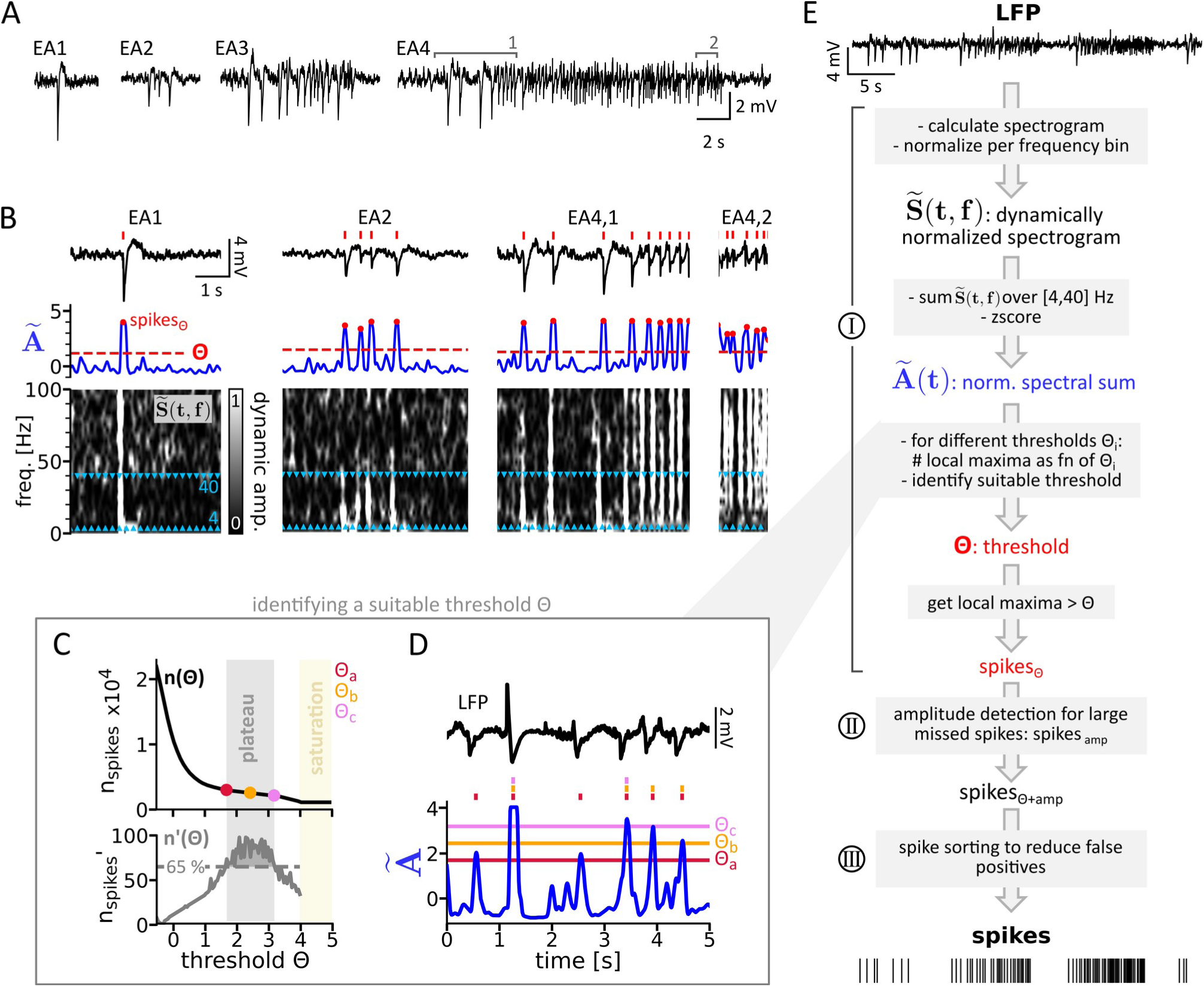
Detecting spikes in the dynamically normalized spectrogram. **(A)** Exemplary EA events recorded from the dorsal hippocampus, close to the kainate injection site. Gray numbers and brackets in EA4 refer to zoomed cutouts shown in B as EA4,1 and EA4,2. **(B)** In the dynamically normalized spectrogram,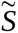 (*t, f*), epileptic spikes manifest as stripes. Summing 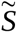(*t, f*) in the 4–40 Hz band and z-scoring results in *Ã* (*t, f*) (blue). Local maxima in *Ã* (*t, f*) above a threshold *Θ* were defined as spikes_Θ_ (red dots, red ticks above the LFP for reference). **(C)** *Top:* The number of spikes *n(Θ)* detected as a function of threshold in one recording session. The colored dots mark the beginning (Θ_a_), middle (Θ_b_) and end (Θ_c_) of the plateau region (gray shade). *Bottom: n’(Θ)*, the slope of *n(Θ)*. The threshold range where *n’(Θ)* was above the 65^th^ percentile, i.e. where the number of spikes was least sensitive to changes of *Θ*, was defined as the plateau region. **(D)** Colored horizontal lines show candidate thresholds Θ_a_–Θ_c_, tick marks indicate spikes detected when applying a certain threshold. **(E)** The workflow followed by PEACOC to transform LFPs into a time series of spikes. Step I (spectral spike detection, illustrated in B–D) is complemented by step II (amplitude detection) to reduce false negatives and by step III (spike sorting) to reduce false positives (see Figure 2).

### From LFP to epileptiform spikes

During larger bursts, spiking often increased in frequency while the amplitude of spikes decreased (EA4 in Figure 1A). During fast and rhythmic spiking, spikes typically had low amplitudes and were not followed by a wave. In these scenarios, neither amplitude thresholds nor template matching would be able to resolve single spikes consistently. Therefore, our detection procedure focusses on the spectral domain (step I, Figure 1B). Spectral spike detection is complemented by an amplitude based detection to enhance sensitivity (step II), and by spike sorting to improve precision (step III, see workflow in Figure 1E).

#### Step I: Spectral spike detection

In PEACOC, the LFP signal of a recording sessions is downsampled to 500 Hz for computational efficiency. Then a spectrogram *S*(*t, f*) is calculated using a fast Fourier transform applied to sliding windows of 256 ms (128 data points), each multiplied by a Hanning window. To reveal fluctuations adequately for all frequencies, the spectrogram is normalized dynamically according to Waldert et al. (2013). In brief, this method normalizes *a* (*t, f*), i. e. the amplitude in temporal bin *t* and frequency bin *f*, relative to all amplitudes in the same frequency bin *a* (:*, f*). The resulting normalized amplitude has values between 0 and 1:

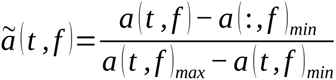

To avoid influences of extreme signal fluctuations *a* (: *, f*)*_min_* and *a* (: *, f*)*_max_*are set to the 5^th^ and 95^th^ percentile of amplitudes, respectively. Values < 0 and > 1 are clipped to keep the value range between 0 and 1.

Epileptiform spikes clearly manifested as stripes in the dynamically normalized spectrograms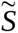(*t, f*)– regardless of whether they occurred isolated, in small bursts or as rhythmic spiking during seizure-like events (Figure 1B). The frequency extent of spectral stripes during spikes varied, but most spikes were accompanied by elevated *ã*(*t, f*) in the 4–40 Hz band. To extract these spectral stripes, amplitudes*ã*(*t, f*) are summed across frequency bins within the 4–40 Hz interval:

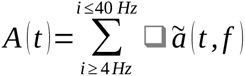

For comparability, *A* (*t*) is z-scored by subtracting its temporal mean and dividing by its standard deviation. The resulting normalized spectral sum*Ã* (*t*)(blue lines in Figure 1B) displayed prominent local maxima coinciding with spikes in the LFP.

Next, a suitable threshold for separating local maximal corresponding to LFP-spikes from local maxima due to noise is identified (Figure 1CD): We define local maxima in *Ã* (*t*) above a threshold Θ as coincident with spikes_Θ_. First, n(Θ), i. e. the number of spikes_Θ_ as a function of Θ, is determined for Θ ranging from − 0.5 to + 6.5 z-scores (resolution: 0.05 z). To avoid double detections, spikes closer than 1/12 s to their predecessors are discarded. The separation between spike maxima and noise maxima manifested as a plateau region in n(Θ): With high Θ, hardly any spikes were detected (Figure 1D). As Θ was lowered towards the plateau region, prominent local maxima corresponding to spikes were detected. Lowering Θ further within the plateau region had little effect on spike count. Once Θ dropped below the plateau region, noise maxima were detected and spike count increased rapidly. Following this observation, a suitable threshold for spike detection should lie within the plateau region of n(Θ). To determine the plateau region, n’(Θ), the derivative of n(Θ) for thresholds below saturation level, is calculated the plateau region is delimited as the shallowest 35 % of n’(Θ). n(Θ) and the plateau region are analyzed for each recording session individually. Spike detection can thus adapt to individual differences in signal-to-noise ratio and spike rate. Note that the plateau region was absent in saline injected mice (*Figure 1—figure supplement 1*), suggesting that n(Θ) could be used to distinguish epileptic from non-epileptic mice.

To determine which part of the plateau region is most suitable to detect spikes, we identified the beginning (Θ_a_), middle (Θ_b_) and end (Θ_c_) of the plateau as candidate thresholds, carried out the remaining detection steps II and III for each, and compared detection performances.

#### Step II: Amplitude-based spike detection boosts sensitivity

While step I correctly identified most spikes, it lead to clear false negatives. Despite high amplitudes and a typical waveform, some spikes were missed because their slopes in the LFP were not steep enough to result in sufficiently high amplitudes at higher frequencies of the 4–40 Hz band (Figure 2A).

**Figure 2:**
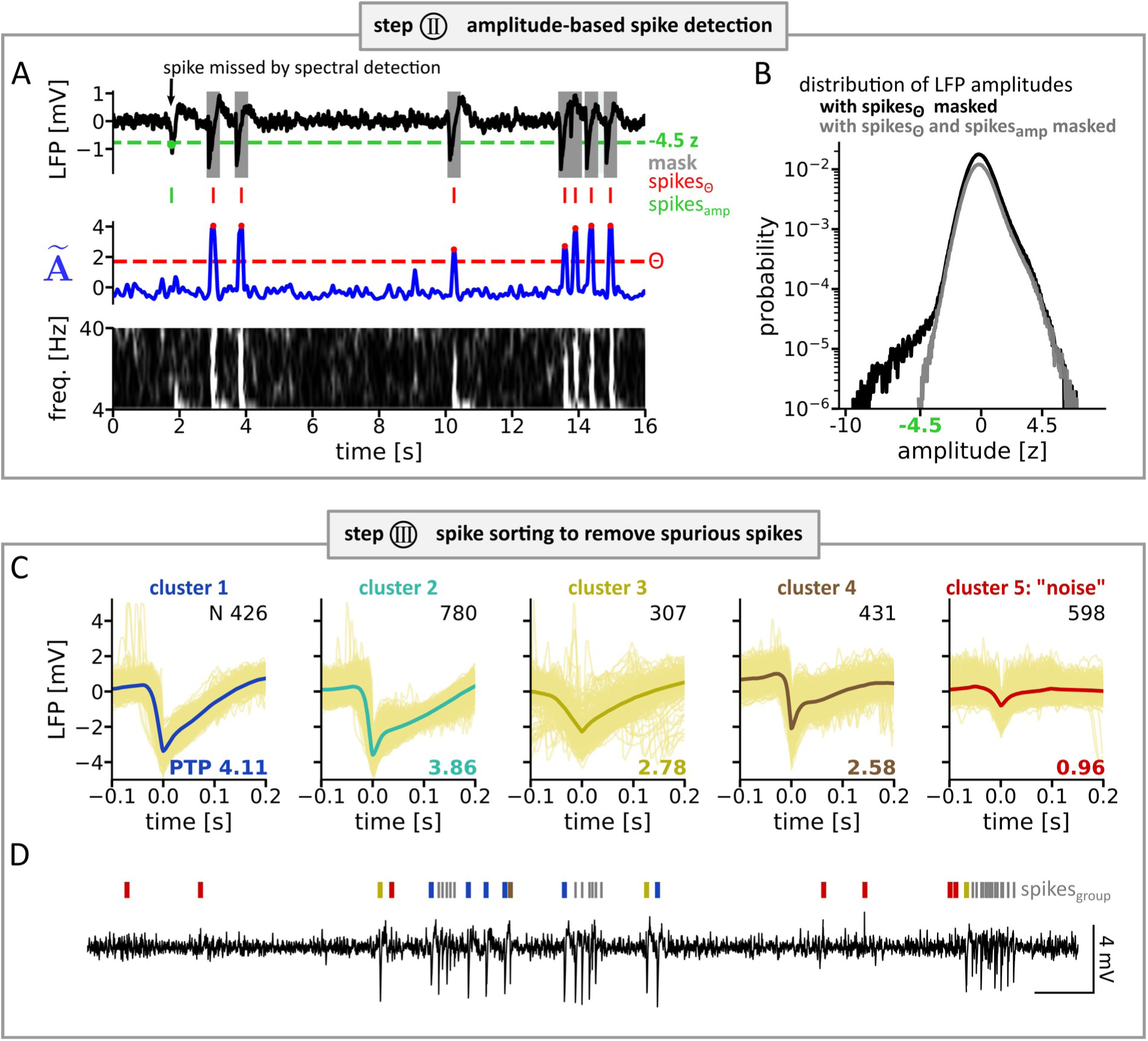
Spike detection based on amplitudes and spike sorting. **(A)** To detect spikes missed by spectral spike detection (step I), spikes_Θ_ (red tick marks) are masked (gray) and an amplitude threshold of − 4.5 z (green line) is applied to the remaining LFP. Threshold crossings are considered as amplitude spikes (spikes_amp_, green tick mark). **(B)** Distribution of a z-scored LFP recording (single session). The amplitude distribution of the LFP with spikes_Θ_ masked (black) displays a shoulder that vanishes when masking both spikes_Θ_ and spikes_amp_ (gray), indicating a homogeneous residual baseline. **(C)** To remove false positives, five spike clusters are identified through spike sorting in each recording session individually (waveforms: thin yellow lines, minima aligned to 0 s, average waveform: thick colored line). Clusters 1 to 5 are sorted for peak-to-peak (PTP) amplitude. The cluster with the lowest PTP amplitude (cluster 5) is considered noise and its spikes are discarded as false positives. **(D)** Exemplary LFP cutout. Tick marks indicate cluster affiliation. Spikes occurring in dense groups (spikes_group_, gray tick marks) are not sorted.

Missed spikes of high amplitude often occurred before a spike burst or in isolation, but we never observed them within dense bursts. Due to this temporal separation between detected and missed spikes, spikes_Θ_ already detected can be masked using 400 ms patches centered on each spike_Θ_ without occluding the missed spikes. After masking, missed spikes manifested as a shoulder at low (for recordings with a negative polarity of spikes, Figure 2B, dark green), high (positive polarity), or low and high (mixed polarity) amplitudes in the amplitude distribution of the remaining LFP. Since shoulders appeared at around 4.5 standard deviations, the time points at which the z-scored LFP crossed a threshold of 4.5 z (− 4.5 for negative spike polarity, + 4.5 for positive polarity and ± 4.5 for mixed polarity) are detected as amplitude spikes (spikes_amp_, Figure 3). Spikes_amp_ accounted for 3.9 % (median across recordings, range: 0.9–13.0 %) of all spikes detected and boosted sensitivity substantially for some recordings and threshold settings (2–68 % higher sensitivity for threshold Θ_c_).

**Figure 3:**
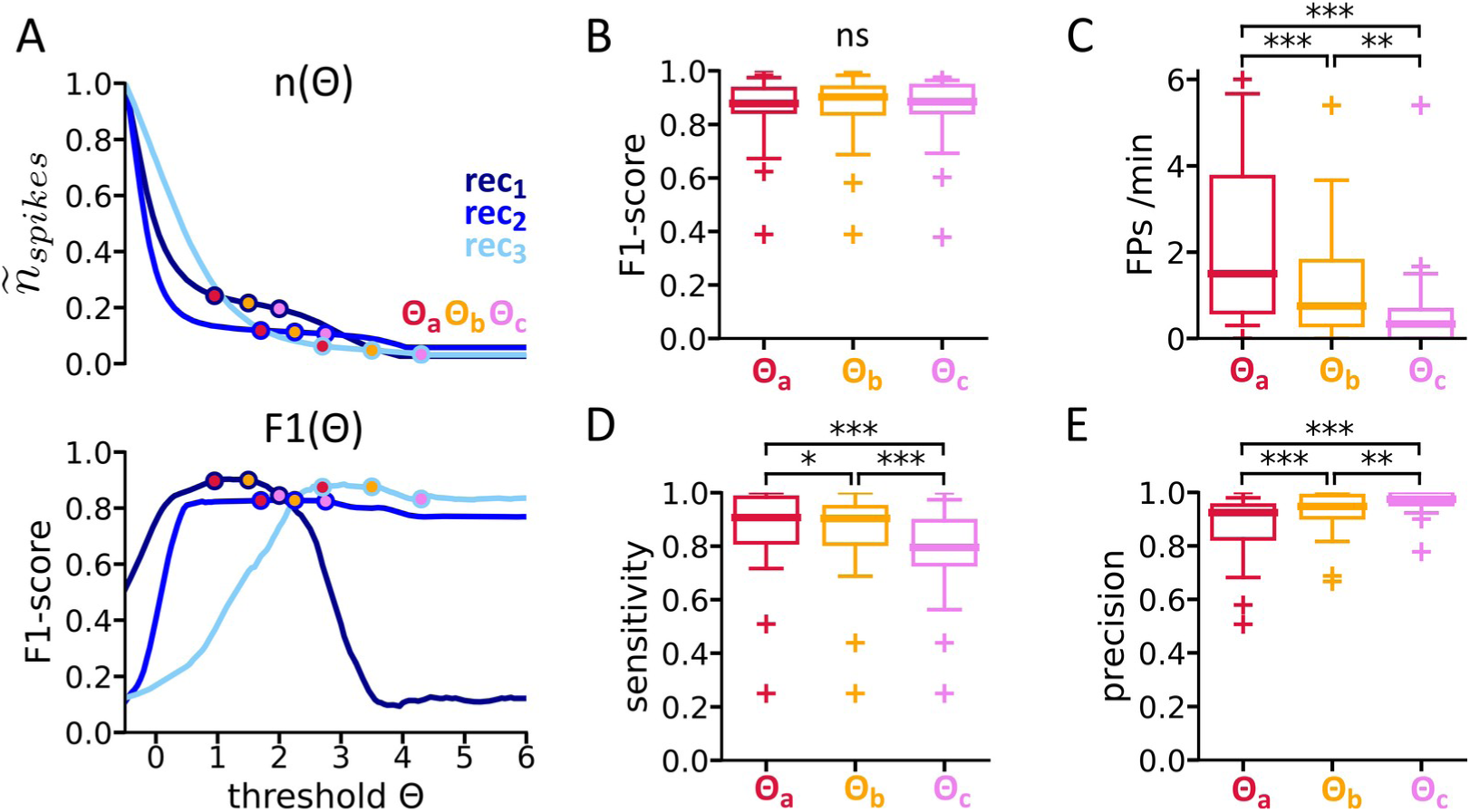
Validation of spike detection. **(A)** *Top: n(Θ)* normalized to maximum = 1 (blue lines) and candidate thresholds Θ_a_–Θ_c_ (colored dots) for example recordings rec_1_–rec_3_. *Bottom:* F1-score (geometric mean of sensitivity and precision) as a function of threshold. **(B–D)** Detections using Θ_a_, Θ_b_ and Θ_c_ do not differ significantly with respect to F1-score (B), but are significantly different in terms of false positive rate (C), sensitivity (D) and precision (E). One sample point per recording (N = 29) and threshold. Within each box, the middle bar indicates the median. Boxes enclose the 25^th^–75^th^ percentiles. Whiskers enclose the 5^th^–95^th^ percentiles. Crosses mark sample points outside the whisker range. Each recording (N = 29) provides one sample point per threshold. Statistics: Kruskal-Wallis test for differences across groups followed by pairwise Wilcoxon signed-rank test with Bonferroni correction; ns p ≥ 0.05, * p < 0.05, ** p < 0.01, *** p< 0.001.

#### Step III: Spike sorting enhances precision

To increase the precision of spike detection, PEACOC next identifies potential false positive (FP) spikes, extracts the corresponding waveforms, and identifies FPs through spike sorting.

FP removal is limited to spikes outside of and at the border of bursts because within bursts spike waveforms could be distorted, overlapping and difficult to separate. Each potential FP is aligned to its minimum (in case of negative spike polarity), respectively its maximum (for positive spike polarity) in the LFP. Windows -100 ms to +200 ms around the minimum respectively maximum are defined as the waveform of the candidate. For recordings of mixed polarity, each candidate is assigned a polarity according to the sign of its most extreme amplitude. Negative and positive candidates are then treated separately in the following steps. After waveform extraction, waveforms are grouped into clusters according to the three principal components that account for the largest variance in the data (Lewicki, 1998). For clustering, a mixture of five Gaussian kernels (Souza et al., 2019) is fitted to these principal components (Python package scikit-learn; RRID: SCR_002577, subclass: GaussianMixture). Finally, spikes belonging to the cluster with the lowest average peak-to-peak amplitude are discarded as FPs (Figure 2CD). Because false detections could mask other potential FPs, the whole procedure is repeated until no new false detections are identified (*Figure 2—figure supplement 1*).

In step III, 5–20 % of spikes (median: 14 %, *Figure 3—figure supplement 1*) were identified as FPs and discarded.

#### Validation: Spike detection is robust against changes in threshold

To validate the detection procedure, we randomly selected 20 s snippets (N = 197) and compared the spikes detected by PEACOC to spikes manually annotated by an expert. F1-scores, i. e. the geometric mean of sensitivity and precision, were maximal for thresholds in the plateau region (Figure 3A). Which of the thresholds Θ_a_–Θ_c_ was used in step I did not have a significant impact on the F1-score of the whole procedure (Figure 3B), indicating that PEACOC’s spike detection performance is robust and that any threshold in the plateau region would be an acceptable choice. However, detections based on Θ_a_–Θ_c_ significantly differed in terms of sensitivity, FP rate and precision (Figure 3C–E). As expected, the lowest threshold Θ_a_ resulted in the highest sensitivity (median: 0.91) but a higher rate of FPs (1.5 /min). Θ_c_ was less susceptible to FPs (0.3 /min) but missed more spikes (median sensitivity: 0.80). The choice of threshold thus allows to emphasize sensitivity (Θ_a_), or precision (Θ_c_), or a compromise between the two (Θ_b_). Note that the differences between the thresholds were larger when just using step I for detection (*Figure 3—figure supplement 1*). Steps II and III had a balancing effect that not only boosted performance but also was essential for the robustness of detection.

The FP rate strongly depended on context (*Figure 3—figure supplement 2*) and was about ten times higher in episodes containing dense spike bursts. The reasons for this are that spike sorting (step III) was only applied to spikes outside or at the border of bursts and that within dense bursts spikes tended to overlap such that spikes could not always be identified unambiguously. This may have led to more false negatives and FPs, which, however, would have little impact on the subsequent classification of bursts due to the already salient spiking characteristics of large bursts.

PEACOC operates with Θ_a_ as a default threshold to emphasize sensitivity and all analysis steps reported here were carried out based on this threshold. Detections using Θ_a_ yielded the best F1-scores in 55 % of recordings, with an F1-score of 0.88, a false positive rate of 2.7 /min, a sensitivity of 0.87 and a precision of 0.89 across all recordings.

### From spike trains to a sequence of classified bursts

Transforming the LFP into a spike train allows PEACOC to delimit EA patterns as spike bursts and to classify these bursts according to spike load.

#### Defining spike bursts based on inter-spike interval statistics

Episodes of high spike rate alternated with predominantly spike-free episodes (Figure 4A). Spike-rich episodes had salient beginnings and ends but varied in spike density and duration. The inter-spike interval (ISI) structure of spiking indicated the existence of discrete bursts events: logarithmic ISI distributions of individual recording sessions were bimodal with one peak at short ISIs followed by a secondary shallower peak or plateau region at longer ISIs (Figure 4B). Following Selinger et al. (2007), we interpreted the first peak as originating from short intra-burst intervals and the second as longer inter-bursts intervals. Accordingly, an ISI in the valley region between the two peaks defined a suitable threshold to delimit bursts. In our recordings, valleys (or sometimes transitions from the first peak to a plateau region) were located at around 2.5 s. Therefore, spikes closer than 2.5 s are grouped into the same burst. In consequence, spikes with intervals > 2.5 s to their neighbors are defined as solitary spikes. False negative spike detections in low amplitude regions occasionally broke up larger bursts. This is corrected by merging bursts closer than 3.5 s. In our dataset, 6.7 % of bursts resulted from such merging of smaller bursts.

**Figure 4:**
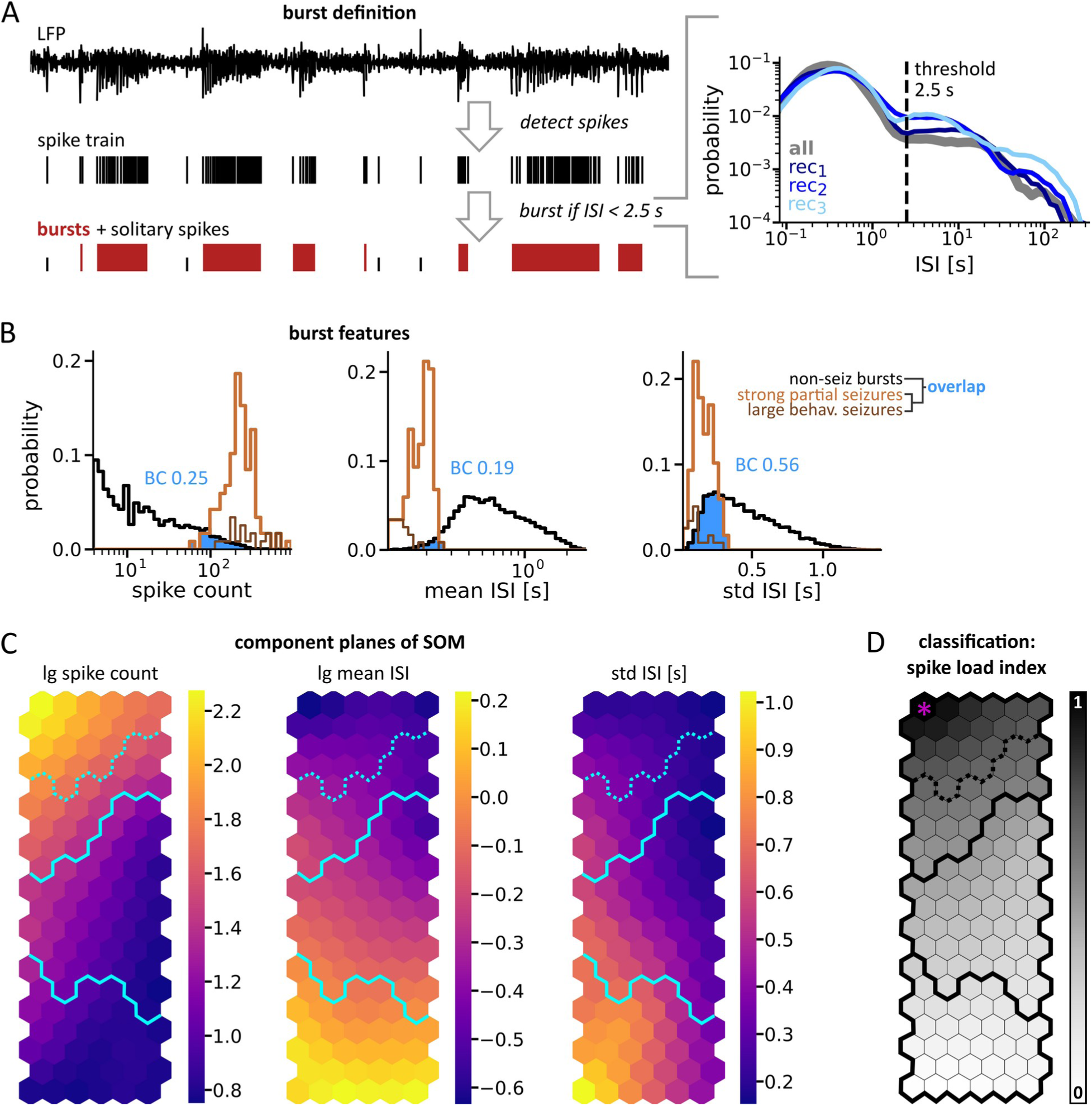
Definition and fine grained classification of spike bursts. **(A) Workflow for burst definition** *Right:* Interspike interval (ISI) distributions of three example recordings (blue colors) and of the whole dataset (gray). Note the valley respectively plateau region at around 2.5 s (dashed line). Spike separated by intervals below threshold are defined as spikes belonging to the same burst. Spikes not belonging to a burst are referred to as solitary spikes. **(B)** Histograms of features extracted from bursts containing at least five spikes (N = 12373). Data from bursts corresponding to visually identified seizures are shown in light brown (strong partial seizures, N = 99) and dark brown (large putatively behavioral seizures, N = 18). Histograms of bursts and seizures (behavioral and partial jointly) were each normalized to an area of 1. The overlap between seizure and non-seizure data (blue area) was quantified with the Bhattacharyya coefficient (BC, blue numbers). **(C)** A self-organizing map (SOM) was trained with bursts from all recordings of datasets A and B (Froriep et al., 2012; Janz et al., 2017). Bursts were represented by three-dimensional feature vectors [2·(lg spike count), 2·(lg mean ISI), std ISI] to obtain prototypical feature vectors (“nodes”, displayed as hexagons) arranged according to similarity. Each component plane depicts the distribution of one of the features with color indicating the respective feature value. Blue lines delimit the broader categories identified in subsequent analyses (Figure 5). **(D)** The spike load index (gray scale) quantifies the similarity of the SOM’s prototype vectors to the prototype vector of the reference node H* (purple asterisk) in feature space. Note the smooth gradient suggesting a continuum. A and B were adapted from Heining et al. (2019).

Since the location of the valley (or the beginning of the plateau) varied between recordings, we tested the impact of different interval thresholds on further analyses. All major findings of Heining et al. (2019) were replicated with interval thresholds ranging from 1 to 5 s (Extended Data Figure 5-3 ibid.), emphasizing the robustness of our approach,.

#### Selecting features for burst classification

Spike bursts varied in numerous features, such as spike rate, duration and regularity (Figures 4B and S5A). Bursts corresponding to manually annotated partial seizures had typically high spike rates and long durations. Yet, other bursts could also have spike rates but be short or could be long with lower spike rates. In some bursts, spiking was periodic, whereas in others the internal spiking pattern appeared less structured or evolved from a loose pattern into rhythmic spiking. Bursts also differed in their average spike amplitude and in amplitude evolution. Especially in longer bursts, spike amplitudes often decreased progressively. This richness of patterns complicates classification.

We aimed to capture the manifold of bursts by quantifying their similarity to electrographic seizures. For this, a suitable feature set should (1) provide extreme values for bursts corresponding to visually identified seizures, (2) be comparable across animals and electrode positions, (3) minimize the number of parameters or thresholds, (4) be non-redundant. Since we considered epileptiform spikes as the smallest unit of EA, we favored spike-based features to express the similarity to seizures. Aside from temporal spike statistics (rate, regularity, etc.), features reflecting amplitudes and signal energy (e. g. line-length), might be interesting since they are popular in seizure detection (Logesparan et al., 2013). Criteria (2) and (3), however, cannot be satisfied easily when using amplitudes. Warranting comparability between mice would require normalization by some amplitude baseline, which is difficult to obtain, especially under epileptic conditions. The temporal evolution of spike rate and amplitude within an event is also commonly used to describe and differentiate EA patterns (Chauvière et al., 2012; Twele et al., 2016a). Evolutionary features, however, are difficult to parametrize and require additional assumptions that would violate criterion (3). We therefore preferred simple spike statistics to evolutionary or amplitude-associated features.

We evaluated the probability distributions of candidate separately for bursts visually identified as seizures and non-seizure bursts (Figure 4B and *Figure 4—figure supplement 1*). When the probability distribution of a feature was heavy tailed, we used the log_10_ transform (abbreviated to lg) of that feature to obtain a more even occupation of the value range. To account for criterion (1), we quantified the degree of overlap between seizure and non-seizure bursts using the Bhattacharyya coefficient. The overlap between seizure and non-seizure bursts was lowest for *lg spike count* and *lg mean ISI*. Jointly these reflect two key aspects: the number of spikes and their rate. Duration is also often used to characterize EA events (Chauvière et al., 2012; Karoly et al., 2017; Maroso et al., 2011a; Riban et al., 2002), yet is fully accounted for by spike count and mean ISI in combination and scores lower separability than these. Since spiking regularity varied across bursts, with often highly periodic spiking during seizures (as also observed by Armstrong et al., 2013 and White et al., 2006), we further included the standard deviation of ISIs within bursts (*std ISI*). Note that for line-length/time the separability between seizure and non-seizure bursts was extremely low. Following these considerations, the features *lg spike count*, *lg mean ISI* and *std ISI* are used in PEACOC’s burst classification. Although these reflect complementary aspects of severity and similarity to seizures, all pairs were significantly correlated (τ_(count, meanISI)_: −0.39, τ_(count, stdISI)_: −0.14, τ_(meanISI, stdISI)_: 0.59; all p_τ_<0.001;). While these features were suitable for our mouse model, other epilepsy models might profit from different feature sets.

#### Classifying bursts according using a self-organizing map

In PEACOC, the feature vectors of each burst event are projected onto a self-organizing map (SOM) to condense the overall distribution of feature combinations into spatially arranged prototype vectors. For each burst event, the best matching prototype vector of the SOM indicates the the burst’s spike load index and a more general category.

The SOM method is particularly well suited for our heuristic classification approach because it visualizes structures of similarity (Kohonen, 2013). A SOM is an unsupervised machine learning algorithm that compresses a set of p-dimensional input feature vectors into a two-dimensional grid of nodes. Each node corresponds to a p-dimensional prototype vector and after training similar prototype vectors are neighbors on the final map.

To train the SOM for EA in kainate-injected mice, we used feature vectors derived from all bursts of at least five spikes (N = 12373, datasets from Froriep et al., 2012 and Janz et al., 2017). Bursts containing fewer than five spikes were not admitted because preliminary tests showed that they would have dominated and distorted the map due to their high incidence (N = 10205) and at times extreme feature estimates. To construct feature vectors, *lg spike count, lg mean ISI* and *std ISI* were extracted from each burst_SOM_ and then z-scored. Since *lg spike count* and *lg mean ISI* scored about two times better separability, these were weighted by a factor of two to enhance their influence. The input to the SOM thus consisted of 12373 bursts *i*, each represented by a feature vector x_i_: *x_i_*=[2 *⋅ z* (*lgspike count _i_*); 2 *⋅ z* (*lgmean ISI _i_*) *; z* (*std ISI _i_*)].

A SOM with 120 nodes provided a sufficiently detailed representation of the input space while preserving neighborhood relationships and keeping the visualization compact (*Figure 4—figure supplement 2*). Using the linear initialization method proposed by Kohonen et al. (2001), we obtained a 6 x 20 (width x height) map architecture and initialized the nodes according to the space spanned by the two largest eigenvectors of the feature covariance matrix. The SOM was trained with the batch method and neighborhood functions provided by Carrasco Kind and Brunner (2014).

To create SOMs for other models of epilepsy, SOM dimensions and feature weights might need to be adjusted, though the general procedure would be the same.

After training, the node at the top left corner held the prototype vector with the highest *spike count*, lowest *mean ISI* and a low *std ISI* (Figure 4C). This node represented bursts with the most severe spiking and we defined it as the reference node H*. The SOM reflected two feature gradients. First, a gradient of *spike count* dominating the top part of the SOM and secondly an opposing gradient of *mean ISI* and *std ISI* dominating the middle and lower part of the SOM. The SOM occupancy reflected differences in occurrence of EA patterns across mice (*Figure 4—figure supplement 3*).

#### The spike load index quantifies the similarity to seizure-like bursts

The best matching unit (BMU) of a burst is defined as the node (“unit”) of the SOM that is closest in Euclidean feature space. The reference node H* was the BMU of most bursts manually identified as putative large behavioral (17 out of 18) or partial seizures (73 out of 99).

We quantified the similarity of node *j* to H* as the Euclidean distance in feature space (*D_j_*) between a node’s prototype vector *h_j_* and H*. Using the transformation *L_j_* =1*− D_j_* / *max* ( *D*_1 : *n*_), we constrained the value range to [0, 1] and defined L_j_ as the *spike load index L* of node *j*. Thus, the spike load index is 1 for H* and 0 for the node furthest away from H* in feature space (*Figure 4E, Figure 4—figure supplement 4*). The spike load index of bursts correlated strongly and linearly with *lg spike count* (τ = 0.72, p_τ_ ≪ 0.001) and *lg mean ISI* (τ = −0.66, p_τ_ ≪ 0.001). In contrast, the relationship between duration and spike load index was weaker (τ = 0.39, pτ ≪ 0.001) and clearly non-linear because short bursts sometimes scored higher spike load indices than longer ones (*Figure 5—figure supplement 1*). Of all candidate features, line-length/time correlated least with the spike load index (τ = 0.18, pτ ≪ 0.001, ibid.).

Visually identified seizures not assigned to H* were exclusively assigned to nodes located in the top row of the SOM and hasd a spike load index close to 1. These nodes and H* were also BMUs for bursts not tagged as seizures (53 % in H*). The spike load index, however, yielded a better separation between seizure and non-seizure bursts than any other candidate variable (Bhattacharyya coefficient = 0.16, compare Figure 4B and *Figure 4—figure supplement 1*).

#### Partitioning the continuum into three categories

In terms of spike load index, the nodes of the SOM formed a continuum: The gradients of the features constituting the spike load index and the gradient of spike load itself were continuous (Figure 4DE, *Figure 4—figure supplement 4*). Moreover, the U-matrix of the SOM, which visualizes distance relationships between neighboring nodes, did not indicate the existence of clearly separable clusters (*Figure 5—figure supplement 2*).

To facilitate further correlation analyses, we partitioned the continuum of spike load into larger categories. We categorized the prototype vectors of the SOM through agglomerative hierarchical clustering (Figure 5AB). The merging distances in the dendrogram increased approximately exponentially at each level of the hierarchy and thus did not suggest specific clusters (Figure 5A, *Figure 5—figure supplement 3*). We cut the dendrogram to yield three categories. These formed contiguous, similarly sized patches on the SOM (Figure 5B) and were arranged along the gradient of the spike load index. Accordingly, we defined these categories as the high-load, medium-load, and low-load category.

**Figure 5:**
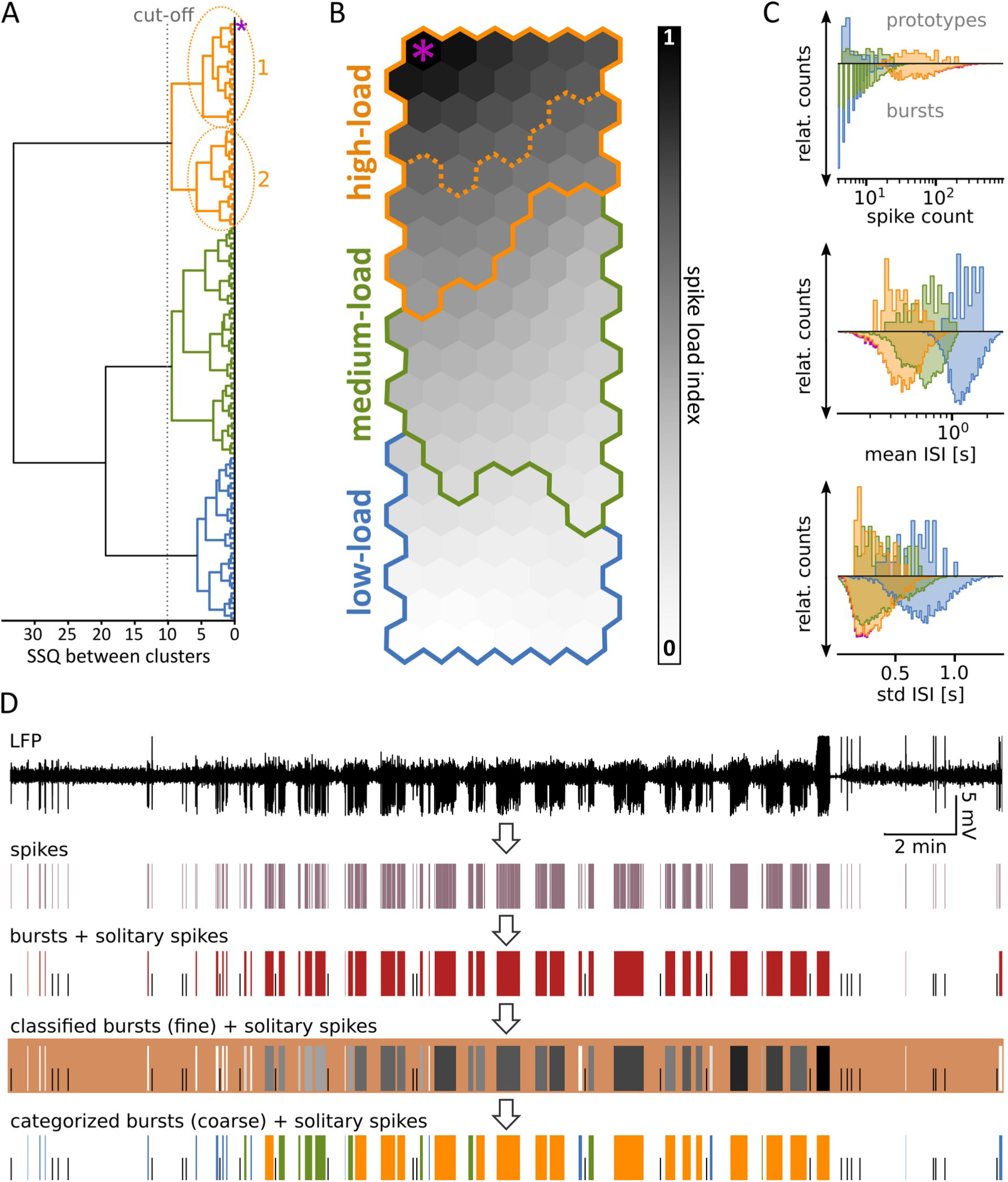
Partitioning the continuum into broader categories. **(A)** Dendrogram derived from merging the SOM’s prototype vectors into clusters. The horizontal axis displays the sum of squares (SSQ) in feature space between merged clusters. The magenta asterisk marks the reference node H*. At the level indicated by the gray dotted line, the dendrogram splits into a high-load, a medium-load, and a low-load branch. The high-load branch could be further subdivided into branches 1 and 2 (dotted line in B). **(B)** SOM with gray levels reflecting the spike load index and colored lines delimiting the categories (branches). **(C)** Distribution of the features used for classification. Relative counts of the SOM’s prototype vectors per category are shown above the gray lines, the distributions of the corresponding burst data below. Bursts inherited their category from their best matching unit (BMU) on the SOM, i.e. from the prototype vector best matching the features of the burst. Histograms were normalized to the same area. Overall, the feature distributions of the bursts were similar to the feature distributions of the SOM’s nodes. **(D)** Overview of workflow. PEACOC detects spikes in the LFP (Figures 1–3) and then groups them into bursts (red boxes, see Figure 4A). Bursts are then classified on a fine scale (spike load index, gray levels) reflecting a continuum (Figure 4B–D) and on a categorial scale (colors, see panel A). A colored background was added to the fourth row to visualize the lighter markers. Panels A and B were adapted from Heining et al. (2019).

For classification, each burst was assigned to its BMU and inherited the category assignment of this BMU. In terms of *lg spike count,* the high-load category was well separable from the low-load and medium-load category, while the distributions of the latter categories overlapped considerably. Conversely, *lg mean ISI*, and to a lesser extent *std ISI*, distinguished the category low-load from the other categories (Figure 5C). Bursts with fewer than five spikes (45 % of all bursts; 10205 of 22578), which had not been admitted to the SOM, were morphological most similar to low-load bursts and were thus assigned to this category.

Beyond the features used for classification, bursts showed complex morphologies and (*Figure 5—figure supplement 4*). We did not observe any sudden qualitative morphological differences at the transitions from high-load to medium-load, and from medium-load to low-load bursts. This further corroborated that burst patterns of EA form a continuum. Event duration and spike amplitudes, which are commonly used for describing and detecting seizures, were only loosely related to the burst categories. Across all categories, spike amplitudes in bursts were of comparable variability and covered similar ranges (Figures S5 and S11). High-load bursts tended to last longer than other bursts, whereas medium-load bursts had overall similar durations to low-load bursts. Apart from generalized seizures, which started with low amplitudes and fast rhythmic activity, most high-load bursts began with a phase of high amplitude spikes at a lower rate. While spike rate could increase during a high-load burst, spike amplitude often decreased. This temporal pattern is similar to the type A bursts observed by Chauvière et al. (2012) in epileptic rats, and the hyperparoxysmal discharges described in our mouse model (Riban et al., 2002; Twele et al., 2016a). On the whole, however, the intra-burst structure of EA in our datasets were variable.

In summary, PEACOC transformes LFP recordings into a succession of solitary spikes and bursts categorized according to spike load (Figure 5D). To test the robustness of our approach, we compared the classifications resulting from alternative thresholds (e. g. for spike detection or burst-definition) and feature weights. When kept within reasonable bounds, these alternative settings produced comparable results.

#### Projecting new datasets on a reference SOM

To obtain reproducible classifications across datasets, we recommend using a SOM derived from a large dataset as a reference or “gold standard”. Therefore, PEACOC itself does not calculate a SOM de novo for each dataset but projects new data onto a pre-defined SOM. Employing a reference SOM has the advantages that it requires less processing time and that datasets that are too small to derive a representative spectrum of EA can be classified relative to a more complete range of EA. To project a dataset in PEACOC, spikes and bursts are detected as described above. Next, a feature vector *y_j_* is extracted from each burst *j*: *y _j_*=[ *lgspikecount, lgmeanISI, stdISI* ]. These *y_j_* are then normalized to the reference dataset (*x*) by subtracting the mean of x (*x*) and dividing by its standard deviation (*σx*). Applying the weights ω of the reference (ω = [2;2;1]) results in a normalized and weighted feature vector 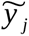 for each burst j: 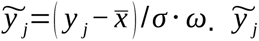 is then assigned to the BMU on the reference SOM to classify burst *j*. We did not encounter any incompatibilities or deficiencies with any novel datasets (Kilias et al., 2018; Paschen et al., 2020; Tulke et al., 2019) projected onto the SOM calculated from the reference dataset (Froriep et al., 2012; Janz et al., 2017). The current SOM accompanying the PEACOC algorithm should thus be suitable to analyze further datasets from the intrahippocampal kainate mouse model. To analyze data from different animal models, we suggest to calculate a new reference SOM from a large enough dataset following the procedures described above.

### Implementation and application

PEACOC’s software architecture and workflow are illustrated in Figure 6. PEACOC can be operated via the operating system’s command-line (tested on Windows, macOS, Linux) or within Python. The data is analyzed per recording session. For a quick overview of EA composition and quality control, PEACOC generates figures in.png or other graphics formats. All results are saved in hdf5 files, which are readily accessible for visualization and subsequent analyses through custom data readers in Matlab and Python. Metadata, such as parameter settings or diagnostic metrics, can be stored in odML (Grewe et al., 2011) or yaml files and rendered in a web-browser or as text. Parameters can be adjusted conveniently, and are saved separately for each analysis run. The modular architecture of the code, allows to adapt parts easily or to incorporate them into other toolboxes. Epileptiform spike detection uses the default threshold Θ_a_, but allows to select Θ_b_ or to adjust the threshold interactively. False positive clusters defined by spike sorting can be selected using a GUI.

**Figure 6:**
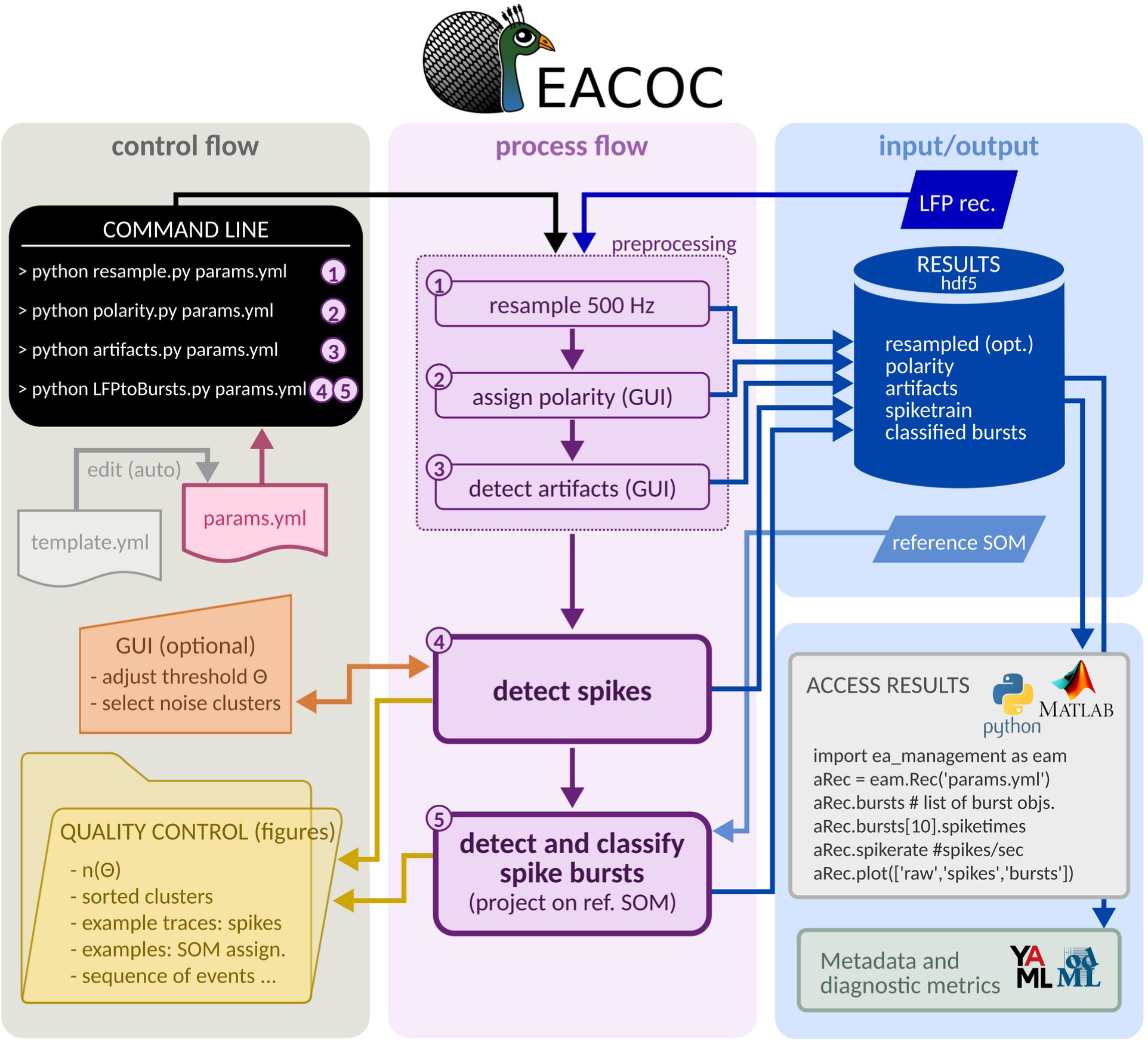
Software architecture and workflow. The PEACOC software transforms a LFP recording into a sequence of classified bursts and solitary spikes. The whole procedure (processes 1–5) can be executed via the command line (black patch). The file param.yml (pink) states from where to retrieve the raw data, where to store data and figures, a reference to default parameters, and any changes to the default parameters. Preprocessing steps 1–3 are optional (artifact times and polarity can also be directly edited and read in .txt format). Spike detection (4) can be executed in interactive mode using a graphical user interface (GUI). Burst detection and classification (5) are fully automated; classification (5) is automated by projecting the detected bursts on a reference SOM (light blue). All main and intermediate results are stored in hdf5 format (blue container). Results, diagnostic metrics and visualizations are accessible through both Python and Matlab (example in gray box). Metadata and diagnostic metrics (e.g. rate of high-load bursts) can be rendered in human readable form (odML or yaml). During steps 4 and 5, diagnostic figures can be saved automatically to allow quality control.

We successfully tested the code on Windows and Linux and macOS platforms (several versions and distributions). At the time of writing, the analysis of a three-hour recording session took consistently less than 10 min on a standard desktop computer. This indicates that our implementation is suitable for real-time analyses. The PEACOC software, an extensive documentation, a tutorial and further examples of application are available at https://github.com/biomicrotechnology/PEACOC.git

### The rate of high-load bursts is anti-correlated to GCD volume

To investigate how the measures of spike load relate to the anatomical pathology of MTLE, we correlated the rates of the different bursts categories against GCD. In line with previous reports, GCD was prominent in the dentate gyrus ipsilaterally to the kainate injection site, but not on the contralateral side (Figure 7A; Heinrich et al., 2006; Riban et al., 2002) and was more pronounced septally, close to the injection site, while it deacreased towards the temporal pole of the hippocampus (Figure 7B; Häussler et al., 2012). Across mice, the rate of high-load bursts was significantly anti-correlated to the volume of GCD (Pearson’s correlation coefficient ρ = −0.67, p_ρ_ = 0.017, Figure 7C), whereas the rates of low-load bursts and medium-load bursts were not significantly correlated to GCD volume (Figure 7DE). Note that the rates of low-load bursts were calculated for background phases, only, i. e. in episodes free of high-load dynamics, to avoid statistical artifacts. Using the overall low-load rate, irrespective of temporal context, would have produced an artificial positive correlation because high-load bursts and associated dynamics automatically block the time available for low-load bursts (compare Heining et al., 2019). Our results show that high-load bursts reflect the degree of pathological degeneration associated with MTLE whereas medium-load and low-load bursts do not.

**Figure 7:**
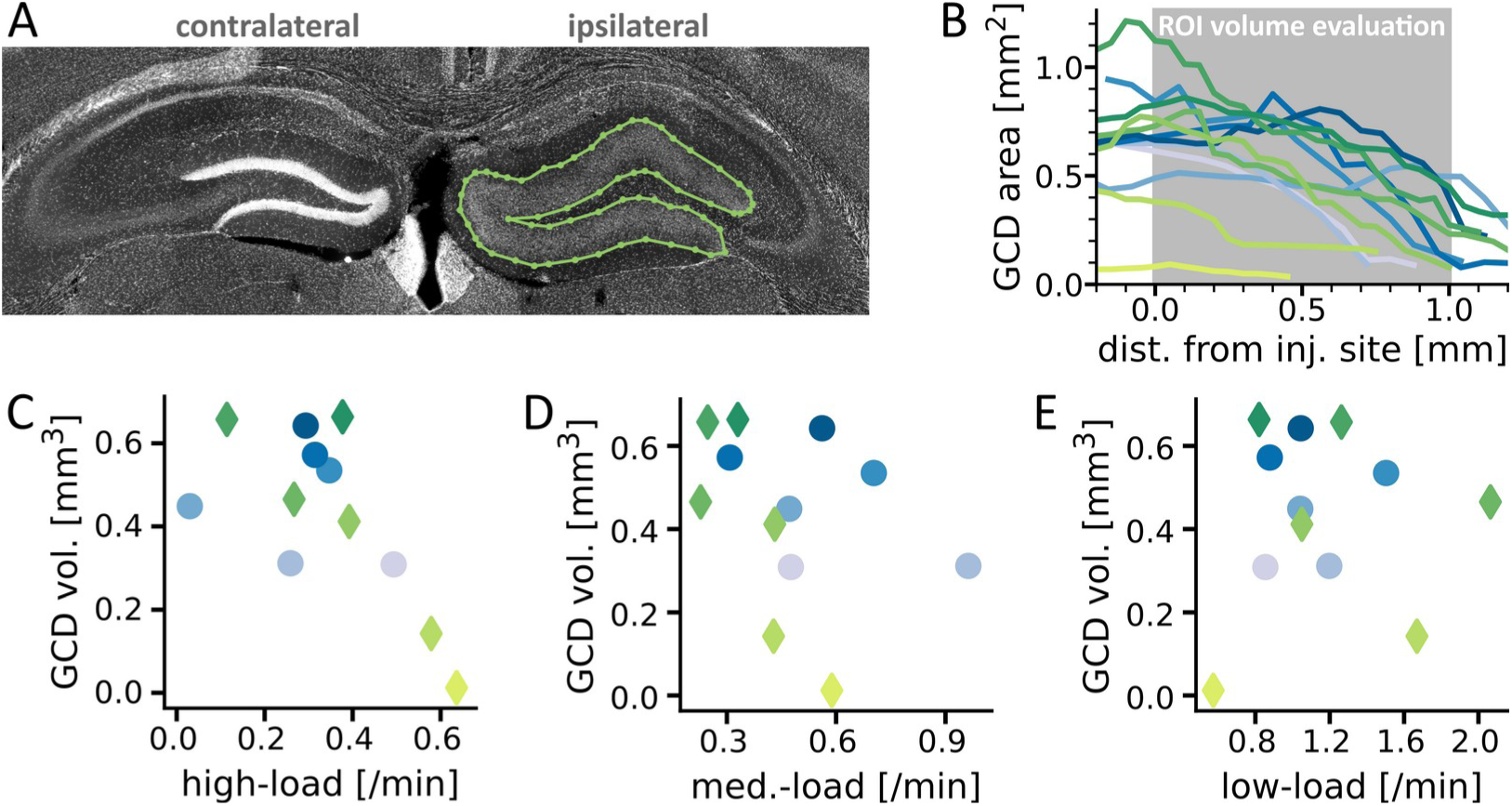
GCD volume is anti-correlated to the rate of high-load bursts. **(A)** Coronal section of a mouse brain showing the dorsal part of the hippocampus. Granule cells are tightly packed on the contralateral side (white band) but dispersed (area encircled in green) on the ipsilateral, kainate injected side. Green dots mark pegging points selected to delimit the area of dispersion. **(B)** Area of GCD as a function of distance from injection site. The volume of GCD was calculated from the injection site to 1 mm further ventrally (gray box). **(C–E)** GCD volume as a function of rate of high-load bursts (C), medium-load bursts (D), and low-load bursts (E). Colors as in B. Panel E shows the background rate of low-load bursts, i. e. the rate of low-load bursts in regions without high-load dynamics. Statistics: GCD vol. vs. high-load rate ρ = −0.67, p = 0.017; GCD vol. vs. medium-load rate ρ = −0.35, p = 0.26; GCD vol. vs. background rate of low-load ρ = 0.08, p = 0.79. Panel B was adapted from Janz et al. (2017).

## Discussion

With PEACOC, we present a software toolbox to automatically detect, delimit, and classify a wide spectrum of EA events based on EA-spike time patterns. PEACOC’s spike detection resolves spikes even within dense bursts and transformed LFP recordings into spike trains. From these, spike bursts are delimited and classified on a fine scale that revealed the continuous character of the EA spectrum with respect to the spike load and into broader categories of high-load, medium-load, and low-load bursts. Correlations with GCD volume and results from other studies using PEACOC (Heining et al., 2019; Janz et al., 2018, 2017; Kilias et al., 2018; Paschen et al., 2020; Tulke et al., 2019) supported the relevance of these burst categories. PEACOC requires minimal user interaction, can derive all critical threshold values automatically from the data, is robust against parameter variations, and thus allows to obtain reproducible results while saving time and resources. Ready access to metadata and diagnostic metrics further enhances PEAOC’s interoperability.

### The benefits of dynamic normalization and adaptive thresholding

Through dynamic normalization (Waldert et al., 2013) we avoided defining a non-epileptic state in epileptic animals as a baseline, which is inherently problematic and may change systematically due to non-epilepsy related factors, such as changes in electrode conductance and scarring of the implanted tissue in long-term recordings.

By dynamically normalizing across the complete recording session, two morphologically identical spikes occurring in two different sessions might appear more or less prominent in the normalized spectrogram depending on the overall spike rate of the recording. To account for this, we did not use a fixed threshold, but derived the spike detection threshold for each recording individually from the characteristic curve n(Θ). In the course of one recording session (1.5–3 h), brain states may change, e. g. from wakefulness to sleep, which could bias spike detection towards brain states accompanied by higher power in the 4–40 Hz band used for detection (see below). To mitigate this effect, the detection procedure could be adapted to each brain state separately as in (Gotman and Wang, 1992, 1991). Alternatively, detection could be performed in extended sliding windows. Preliminary analyses indicate that operating the spike detection in sliding windows of approximately 20 min results in stable detection performance.

### Spike sorting may be useful beyond detecting false-positives

Amplitudes and morphology of spikes depend on brain state, noise amplitude, electrode configuration and, most importantly, the generating network (Demont-Guignard et al., 2009; White et al., 2006; Wilson and Emerson, 2002). One prominent distinction has been made between spikes with and without a wave component (type 1 respectively type 2, Buzsáki et al., 1991). Type 1 spikes were interpreted as generated by a large population encompassing excitatory and inhibitory neurons and type 2 spikes as generated by a more local, exclusively excitatory population (Chauvière et al., 2012). There is strong evidence that the LFP signature of spikes also provides insights into the maturation of epilepsy (Chauvière et al., 2012; Huneau et al., 2013) and their immediate effect as pro- or anti-epileptic (Chauvière et al., 2012; de Curtis and Avoli, 2016). In clinical contexts, automatic sorting of spikes according to their morphology has been used to localize seizure onset zones and to predict surgical outcomes (Coutin-Churchman et al., 2012; Pedreira et al., 2014). Though we used spike sorting merely to reject false positives, differentiating between types of spikes could help to understand EA dynamics in the intrahippocampal kainate mouse model.

### Dentate spikes and sharp waves

In some cases, PEACOC may detect dentate spikes and sharp waves that reflect synchronous population activity during episodes of large irregular activity accompanying slow-wave sleep, resting or consummatory behaviors (Headley et al., 2017). Spectrograms of control data showed prominent stripes during dentate spikes, similar to the stripes accompanying epileptiform spikes. Sharp waves, on the other hand, have a lower amplitude, typically last longer than epileptiform spikes (Buzsáki et al., 2003) and showed low power 4–40 Hz in the spectrograms of control animals. It has been hypothesized that epileptiform spikes are an augmented – and sometimes reverberating (in a burst) – version of sharp waves (Buzsáki, 2015). Considering the mechanistic differences between sharp waves and dentate spikes, Bragin et al. (1995) proposed that the epileptiform type 1 and type 2 spikes are pathological versions of dentate spikes respectively sharp waves.

The similarity between epileptiform spikes and physiological spikes makes it difficult to differentiate them from one another. Our algorithm might therefore systematically detect more false positives during periods with sharp waves and dentate spikes. In healthy rats, however, the rates of dentate spikes (0.2–0.9 /min during slow wave sleep, Headley et al., 2017) and sharp waves (0.2–0.6 /min, ibid.) are far lower than the spike detection rate of PEACOC (10–71 /min) in epileptic mice. Moreover, the physiological patterns of sharp waves and dentate spikes are more irregular compared to the patterns of epileptiform spikes. Since our recordings were obtained during periods of activity, the contribution of falsely detected physiological spikes to overall detections is likely negligible. Conversely, if physiological spikes arise from related network mechanisms, it might be desirable to include them with epileptiform spikes.

### Validation in the absence of a ground truth

Lacking a gold standard, the difficulty to decide whether a deflection in the LFP is indeed a spike leads to discrepancies between experts annotating spikes, especially when noisy (Gaspard et al., 2014; Wilson and Emerson, 2002). While the spike detection of PEACOC was fully validated by one expert only, routinely reviewed randomly selected snippets of LFP marked with the spikes detected by users in other laboratories did not indicate bias and or major mistakes, such as missing indisputable spikes or detecting clear false positives.

### Comparison to other spike detection algorithms

PEACOC detects most spikes in the spectral domain and the amplitude-based spike detection intercepts a small fraction of missed large amplitude spikes. Detecting events solely through an amplitude threshold has been common practice when the desired spikes have large amplitudes. For the kainate mouse model of MTLE, Armstrong et al. (2013) and Krook-Magnuson et al. (2013) used a manually adjusted amplitude threshold to identify regular spiking at the onset of seizure-like events. Such amplitude thresholds are, however, not suitable to detect spikes that occur late within bursts when amplitudes decrease (Krook-Magnuson et al., 2013). Similarly, template matching is popular for spike detection (El-Gohary et al., 2008; Karoly et al., 2016), but fails when waveforms are highly variable or cannot be clearly delimited – e.g. within bursts. Huneau et al. (2013) detected spikes in kainate-injected mice as transient increases of power in the 30–70 Hz band, yet observed that this type of detection was insensitive to spike bursts. In rats systemically injected with kainate, White et al. (2006) calculated the maximum slope in 16 ms sliding windows and defined spikes as outliers of an expected distribution. Unfortunately, the expected distribution is sensitive to the spike rate, which can be very high in our mouse model. Moreover, the wide range of spike morphologies and transients – especially within bursts – is problematic for a slope measure. Bergstrom et al. (2013) used wavelet decomposition corresponding to 0–25 Hz to detect EA in mice with systemic kainate injection. They distinguished episodes of abnormal activity, spiking, and electrographic seizures, but could not resolve spikes individually in these contexts.

### A continuum approach for detecting and classifying spike bursts

EA is often addressed as either individual, “interictal” spikes on the one extreme or seizures (i. e. ictal activity) on the other, with a gap between the two concepts (e. g. Lévesque et al., 2016). Yet, in both, animal models and patients, a wide range of EA patterns can be found and it is not clear whether bursts of spikes, subclinical seizures, and partial seizures can be distinguished at all (D’Ambrosio and Miller, 2010; de Curtis et al., 2012; Gotman, 2011; Sperling and O’Connor, 1990; Twele et al., 2016a; Zangaladze et al., 2008b). PEACOC fundamentally treats EA as a continuum but also derives broader categorizations from this continuum, which are useful when further analyses (e. g. correlation statistics) require a higher level of generalization.

### Comparison to other classification algorithms

Beyond spike and burst characterization, PEACOC is intended to map the continuum of EA.

Supervised machine learning methods are commonly used to classify temporal windows as seizure or non-seizure periods, (Guo et al., 2010; Nigam and Graupe, 2004) (Gardner et al., 2006) but are not ideal to map a continuum. Similar to our study, Meier et al. (2008) observed that SOMs could well separate seizure epochs from non-seizure epochs, map the similarity structure of related seizure types, and distinguish seizures from interictal activity. However, as their goal was real-time seizure detection, they fed the features developed with the SOM to a support vector machine with binary output and did not further pursue the sub-classifications suggested by their SOM.

In contrast to classifications based on supervised machine learning, other approaches extract features and then directly employ one or multiple manually set thresholds in combination. These thresholds are either applied automatically (Armstrong et al., 2013; Bergstrom et al., 2013; Krook-Magnuson et al., 2013), or enforced by an expert annotating the data (Maroso et al., 2011b; Meier et al., 2007; Muhammad et al., 2018; Riban et al., 2002; Twele et al., 2016a). As for the supervised machine learning methods, the problem with such expert-based thresholding approaches is that they focus on detecting a single type of event which is presumed to exist a priori and whose definition is tailored towards the bias of an expert. Armstrong et al. (2013), for instance, developed a successful online event detector for our mouse model (Krook-Magnuson et al., 2013; Zeidler et al., 2018) that reportedly can be tuned to detect a wide range of EA event types. This detector extracts several features, which an experimenter then selects, thresholds, and combines in a series of AND/OR combinations to yield a binary detection decision. To obtain the desired performance of seizure detection, these parameters need to be tuned individually for each animal. The binary output and substantial degree of individualized tuning involved renders these kinds of methods unfeasible for our purposes.

The features used by PEACOC are entirely based on spike statistics. Many seizure detection algorithms employ measures related to signal amplitude, such as energy (Gardner et al., 2006) and line-length (Esteller et al., 2001), which are sometimes used in combination with a wavelet transform to focus these measures on particularly seizure-relevant aspects of the signal (e.g. Bergstrom et al., 2013). While for instance line-length has been successfully used for seizure onset detection when operated on windows of fixed duration (Logesparan et al., 2013), we found this feature inadequate to further classify the already detected and delimited burst events. The reason for this may be the tight association between line-length and amplitudes, which can dramatically decrease in the course of severe bursting.

### High-load bursts resemble electrographic seizures

High-load bursts strongly resembled the events others manually annotated as electrographic seizures. Studies of our animal model commonly distinguish between two sub-types of electrographic seizures: high-voltage sharp waves (HVSWs) and hippocampal paroxysmal discharges (HPDs, both comprehensively reviewed in Twele et al., 2016). HVSWs show a characteristic succession of monomorphic high-amplitude spikes. HPDs, on the other hand, typically begin with HVSW-like spiking that then evolves into spikes of lower amplitudes at higher frequencies. To distinguish HPDs and HVSWs from each other and from interictal activity, experts annotating the LFP are typically instructed to consider some thresholds of duration, amplitude, spike rate, and, sometimes, minimum inter-event interval. The minimum of inter-seizure interval of 3 s suggested by Twele et al. (2016) based on experience happens to lie between the threshold interval of 2.5 s we employed to define bursts and our minimum inter-burst interval of 3.5 s.

Although the various descriptions and definitions of HPDs and HVSWs appeared to be exclusively classified as high-load burst by our method, we observed a continuum in morphology between HVSW-like and HPD-like phenomena and some events with paroxysmal-like spiking did clearly not fit into the molds of either HVSWs and HPDs.

### Correlations with markers of hippocampal sclerosis confirm the relevance load-based categorizations

The anti-correlation between the rate of high-load bursts and overall GCD volume suggests that EA in the category high-load burst is more frequent in less damaged brain tissue. As we showed in Janz et al. (2017), the volume of GCD in turn strongly correlates with the degree of network restructuring and cell death in the sclerotic hippocampus. In the same study, we demonstrated an anti-correlation between seizure-like bursts and a compound variable reflecting cell death and restructuring for dataset A of the present study. The anti-correlation between GCD and high-load bursts we report here for a larger dataset (dataset A + dataset B from Froriep et al., 2012) thus corroborates the idea that more disrupted networks tend to generate fewer severe EA events. While this might suggest that anatomically more affected hippocampi produce fewer high-load bursts in total, this is not conclusive since we evaluated EA only in the dorsal part of the hippocampus, close to the kainate injection site, where cell death, loss of neurogenesis and restructuring is most pronounced (Häussler et al., 2012a; Marx et al., 2013). Yet, the burden of EA is much stronger in the more ventrally located intermediate zone and in the non-sclerotic contralateral hippocampus (Häussler et al., 2012a; Janz et al., 2018). It is further possible that the septal, sclerotic part of the hippocampus does not initiate most EA but that instead EA is relayed to it from non-sclerotic parts. This propagation could be increasingly impaired with progressing sclerosis, leading to EA with lower spike load in the dorsal dentate gyrus. To better understand where epileptiform spikes and bursts are generated, how they propagate, and in what way they relate to anatomical changes, analysis of EA at multiple locations along the septo-temporal axis of both hippocampi will be needed.

### Areas of application

PEACOC was designed using the intrahippocampal kainate mouse model, which displays high rates of epileptiform spiking (Riban et al., 2002; Twele et al., 2016a). A high spike rate was required to obtain a characteristic plateau region, which was in turn necessary to determine automatically a threshold for spectral spike detection. Yet, even without a plateau it might still be possible to retrieve spikes through selecting a high threshold, e. g. at four standard deviations of *Ã* (*t*). Future studies may address whether the original approach or such a modified approach could be used to detect spikes in female mice, where strong spike bursts are rare (Twele et al., 2016b), and in other mouse strains, where the composition of EA may differ (ibid). Preliminary tests indicate that both the detection and classification algorithm are suitable for a novel genetic mouse model of epilepsy and for chronically kindled rats.

We used two datasets (Froriep et al., 2012; Janz et al., 2017) to derive a reference SOM on which PEACOC projected data from other studies to obtain comparable classifications. The success and validity of using the projection method hinges on a comprehensive and valid representation of EA patterns in the reference SOM. Thus far, employing the default spike detection combined with the SOM projection method resulted in a plausible mapping of EA patterns from other datasets (Kilias et al., 2018; Tulke et al., 2019), even when lower doses of kainate had been used to induce epilepsy (Paschen et al., 2020). Future studies should, however, seek to systematically validate the projection method (e. g. by comparing quantization errors and topographic errors, see *Figure 4—figure supplement 2*) and if needed create a more comprehensive mapping of the EA landscape through a SOM that is based on more datasets. Analyzing data of animal models with a different interspike interval structure or different types of epilepsy or EA, might require a customized burst definition, feature selection, and specialized SOMs.

If and how our approach can be applied to evaluate EA data from patients is not clear. As White et al. (2006) argue, electrophysiological signals are more complex in the human brain, and the pathology of human MTLE is more diverse compared to animal models of epilepsy.

### Outlook

Preliminary tests suggest that PEACOC’s spike detection and burst-classification can be adapted for online detection and classification. PEACOC could thus become part of a closed-loop system to monitor and reduce seizure susceptibility, e. g. by detecting phases of high susceptibility (indicated by low rates of low-load bursts, Heining et al., 2019) in real-time, upon which low-frequency stimulation (Paschen et al., 2020) – or possibly stimulation with artificial low-load bursts – would be triggered to prevent high-load bursting.

## Materials and Methods

Data presented in this study and used for the development of PEACOC originates from two datasets obtained for previous publications: set A from Janz et al. (2017) and set B from Froriep et al. (2012). Experimental procedures are described in these studies in detail and briefly below.

### Animals and kainate injection

Experiments were conducted with adult male C57B1/6N mice, housed at room temperature in a 12 h light/dark cycle with food and water ad libitum. All animal procedures were in accordance with the German Animal Welfare Act and were approved by the ethics committee of the regional council in Freiburg, Germany, all procedures for animals from set A were in accordance with the guidelines of the European Community’s Council Directive of 22 September 2010 (2010/63/EU). To induce chronic epilepsy, mice were anesthetized by intraperitoneal injection of ketamine hydrochloride (100 mg/kg body weight) and xylazine (5 mg/kg, atropine 0.1 mg/kg), mounted on a stereotactic frame, and injected with kainate once (50 nL of 20 mM kainate solution in 0.9 % saline solution) into the right dorsal hippocampus as described in Heinrich et al. (2006). After injection, all mice used in this study showed behavioral signs of SE (compare Riban et al., 2002).

### Electrophysiological recordings

At least two weeks after the kainate injection, when mice had reached the chronic stage of epilepsy (Heinrich et al., 2006; Riban et al., 2002), recording electrodes were implanted. Mice were implanted at various locations within the ipsilateral and contralateral hippocampus but the data used in this study originates from electrodes placed in the granule cell layer or molecular layer of the dorsal right hippocampus, close to the injection site. After recovery from surgery, mice were recorded for 1–3 h every day (set A) or on alternating days (set B). The recording period lasted for five days (set A) or up to three weeks (set B). LFPs were amplified (x1000), bandpass-filtered (1 Hz to 5 kHz) and sampled at 10 kHz. During the recordings, mice of set B were monitored with a video camera. We obtained 4–10 sessions per mouse for further analyses.

### Histology

After experiments, mice were anesthetized and transcardially perfused with 4 % para-formaldehyde solution. To verify electrode position and assess histopathological changes, brains were extracted, sliced with a vibratome, and stained (set A: DAPI, set B: Nissl). GCD was quantified in detail for all mice in set A and a subset of set B. Since the granule cell layer is known to widen progressively in the kainate model (Suzuki et al., 1995), we limited the detailed quantification of GCD to those six mice of set B that had been perfused after a similar time following kainate injection (29–32 days) as the mice of set A (36 days). The area of GCD was quantified by pegging the extent of the granule cell layer. Next, the areas between slices were linearly interpolated (step-size: 1 μm) and the GCD volume was calculating between the injection site to 1 mm further temporally because data was available in this interval for most mice.

### Validation

#### Spike detection

We compared spikes detected by our algorithm to spikes selected by a human expert. In total, the expert marked 1550 spikes in 197 snippets, each lasting 20 s. Snippets were randomly selected from 29 recordings (4–10 snippets per recording). For a context dependent evaluation of detection performance (*Figure 3—figure supplement 2*) each snippet was manually labeled as either free (no spikes, N = 65), sparse (a few loosely spaced spikes or small bursts, N = 98) or burst (dense bursts of spikes, N = 33). To quantify detection performance, automatically detected spikes (S_auto_) were matched to visually annotated spikes (S_visual_). Iterating successively through all S_visual_ within a snippet, the S_auto_ closest to a S_visual_ was identified as a true positive (TP) if it occurred within a ± 150 ms window of S_visual_. Only a single S_auto_ could be matched to each S_visual_. Any S_auto_ not matched to a S_visual_ was defined as a false positive (FP). If there was no S_auto_ within ± 150 ms of a S_visual_, this S_visual_ was registered as a false negative (FN). Detection performance was summarized as follows: (1) sensitivity, i. e. the fraction of visually selected spikes that were also detected by the algorithm, (2) precision, i. e. the fraction of automatically detected spikes that had also been selected visually, (3) F1-score, i. e. the geometric mean of sensitivity and precision, and (4) false positives per minute (FP /min).

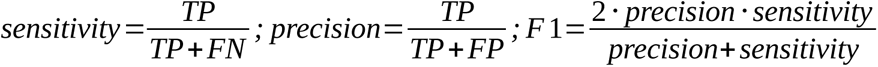

### Strong partial and large behavioral seizures

To obtain a reference for the severity of bursts, a human expert marked putative seizures in all recording sessions. Putative large behavioral seizures were readily identified by their characteristic morphology (fast rhythmic spiking with characteristic amplitude modulation followed by depression of the LFP) and strongly synchronous activity in another electrode channel (set A: contralateral septal hippocampus; set B: medial entorhinal cortex). The definition of strong partial seizures was more uncertain and subjective. The expert tagged events that appeared severe, such as comparatively long, intense bursts of epileptiform spikes.

### Software and code

We developed all custom software, conducted the analyses, and created the graphs presented in this study with Python version 3.5 (Python Software Foundation; RRID:SCR_008394). The code, its documentation and a tutorial are available at https://github.com/biomicrotechnology/PEACOC.git.

### Statistics

Standard statistical tests were conducted with the Python 3 module scipy.stats (Jones et al., 2001). If not stated otherwise, Kendall’s τ (corresponding p-value: p_τ_), which is a rank-based correlation coefficient that does not require normally distributed data, was used. Differences between groups of samples were first tested using the Kruskall-Wallis test. If there was a significant (p<0.05) difference between groups, pairs of groups were compared using the Wilcoxon signed-rank test and pairwise comparisons were corrected with the Bonferroni method. If not stated otherwise, values and ranges are given as the median and the [10^th^, 90^th^] percentile interval. To quantify the overlap between two distributions, their histograms *p* and *q* were calculated with identical *m* bins and the Bhattacharyya coefficient *BC* was determined according to Comaniciu et al. (2000) as

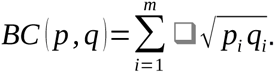

For correlating event categories against GCD (Figure 7), the overall rates of events categories were calculated as duration weighted averages across recording sessions from a mouse.

### Data analysis and pre-processing

To reduce the transient effects of the brief isoflurane anesthesia applied when connecting the headstage with the recording setup, the first 10 min of each session were discarded. Before detecting spikes, artifacts were masked and the dominating polarity of spikes (polarity of the largest peak) was determined in each recording.

#### Masking artifacts

Experts inspected LFP recordings visually aided by a custom algorithm that suggested candidate artifacts. The algorithm detected (1) large amplitude deflections (≥ 4 standard deviations) accompanied by long intervals in which the zero-line was not crossed (≥ 0.5 s), and (2) possible saturation artifacts. Experts could dismiss these potential artifacts and could mark additional regions not detected by the algorithm. For set B they could also consult video recordings. All artifacts were masked before further analyses.

#### Assignment of polarity

Spike polarity depends on the electrode location with respect to the source of a spike. If both positive and negative spikes were common, mixed polarity was assigned. Of all recordings in this study, 74.3 % had negative, 1.9 % positive, and 23.8 % mixed polarity. Assigning polarity allows to apply a polarity-specific amplitude threshold, to align spikes to their peak, and to treat positive and negative spikes separately during spike sorting (if polarity is mixed).

#### Selection of candiate events for spike sorting

Detected spikes were considered potential FPs, if they had fewer than 4 spikes during the preceeding 3 s AND were surrounded by fewer than 5 spikes in a window of 4 s AND were followed by fewer than 5 spikes in a window of 2 s.

#### Clustering of prototype vectors

Using Ward’s method, the closest pairs of clusters, i. e. the pair whose merging would result in the lowest increase of overall within-cluster variance, were merged successively (Ward, 1963). Euclidean distance was used to determine the within-cluster variance.

