## supplementary figures and captions for "PEACOC - Detecting and classifying a wide range of epileptiform activity patterns in rodents"

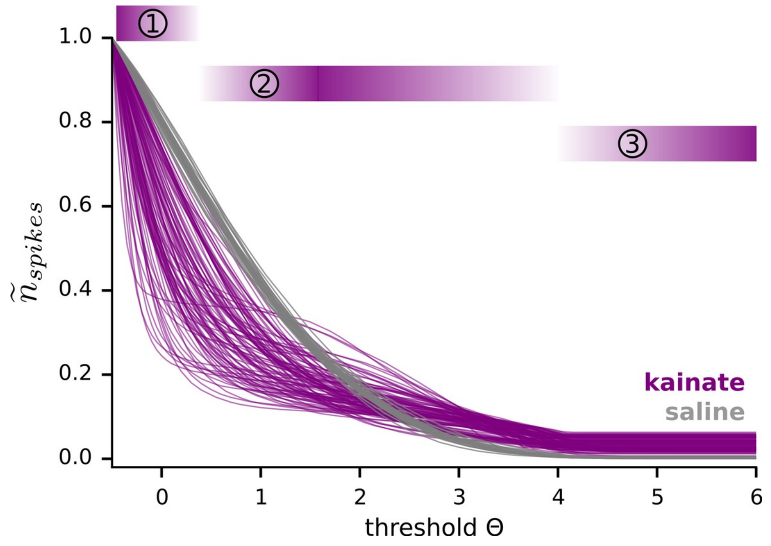


**Figure 1—figure supplement 1: Epileptic and control mice differ in terms of n(Θ)**

The number of spikes detected as a function of threshold Θ for recording sessions from kainate (pink, N = 105 sessions) and saline injected mice (gray, N = 27). For visualization n(Θ) is normalized to a maximal value of 1. Note the difference between kainate and saline treated mice in terms of (1) initial decay, (2) the existence of a plateau region, and (3) saturation levels. In epileptic mice, the exponential decay at low thresholds was much steeper and then settled into a clear plateau for intermediate thresholds. Conversely, control mice maintained an exponential decay in n(Θ) throughout a wide range of intermediate thresholds. At high thresholds, n(Θ) reached saturation levels in epileptic mice, but decayed to zero in control mice.

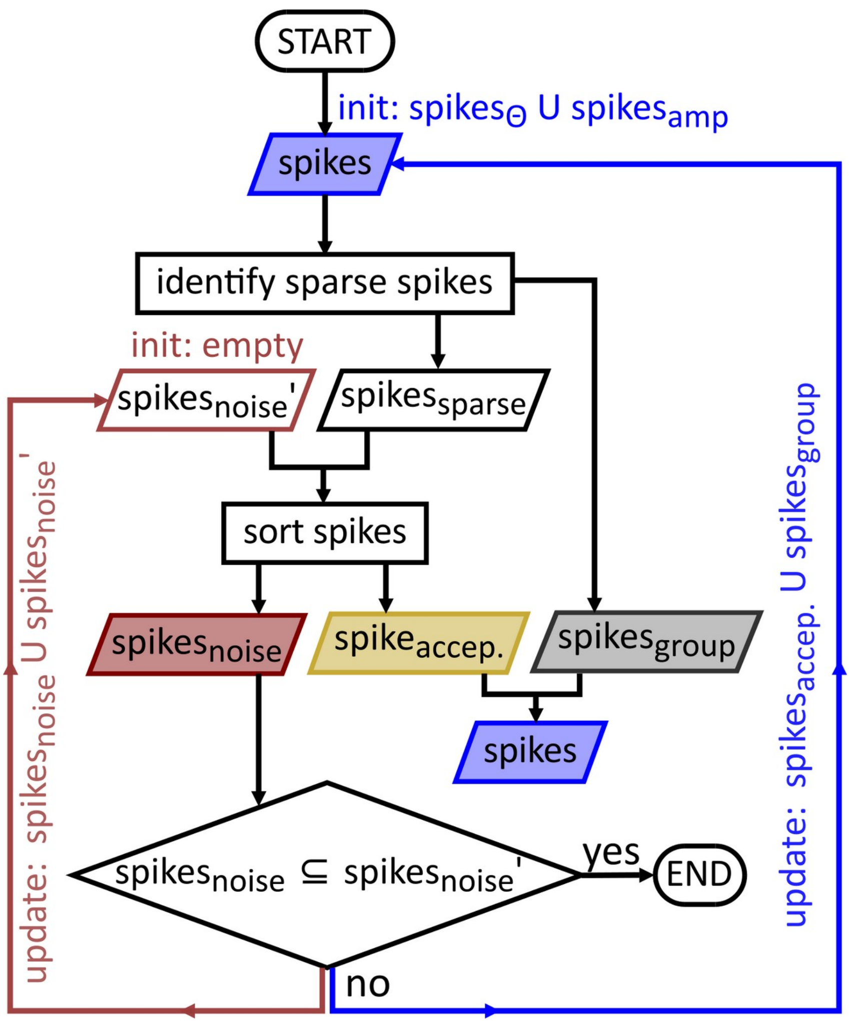


**Figure 2—figure supplement 1: Recursive identification of false detections**

Initially, all spikes (top blue box) are split into two sets: those outside or at the border of spike bursts (spikes$_{sparse}$) and those within bursts (spikes$_{group}$, gray). Spike sorting then determines which spikes$_{sparse}$ are accepted (spikes$_{accep.}$, yellow) and which are considered false detections (spikes$_{noise}$, red). The procedure of identifying sparse spikes is repeated for all spikes except spikes$_{noise}$, i. e. the combined set of spikes$_{group}$ and spikes$_{accep.}$(*spikes*, bottom blue box). Spikes$_{noise}$, on the other hand, directly enter the next round of spike sorting and are stored (spikes$_{noise}$') and masked for the identification of new spikes$_{sparse}$. Repeating the identification of spikes$_{sparse}$ without spikes$_{noise}$, thus prevents spikes$_{noise}$ from masking other false detections as spikes$_{group}$. This procedure is halted once no new spikes$_{noise}$ are identified.

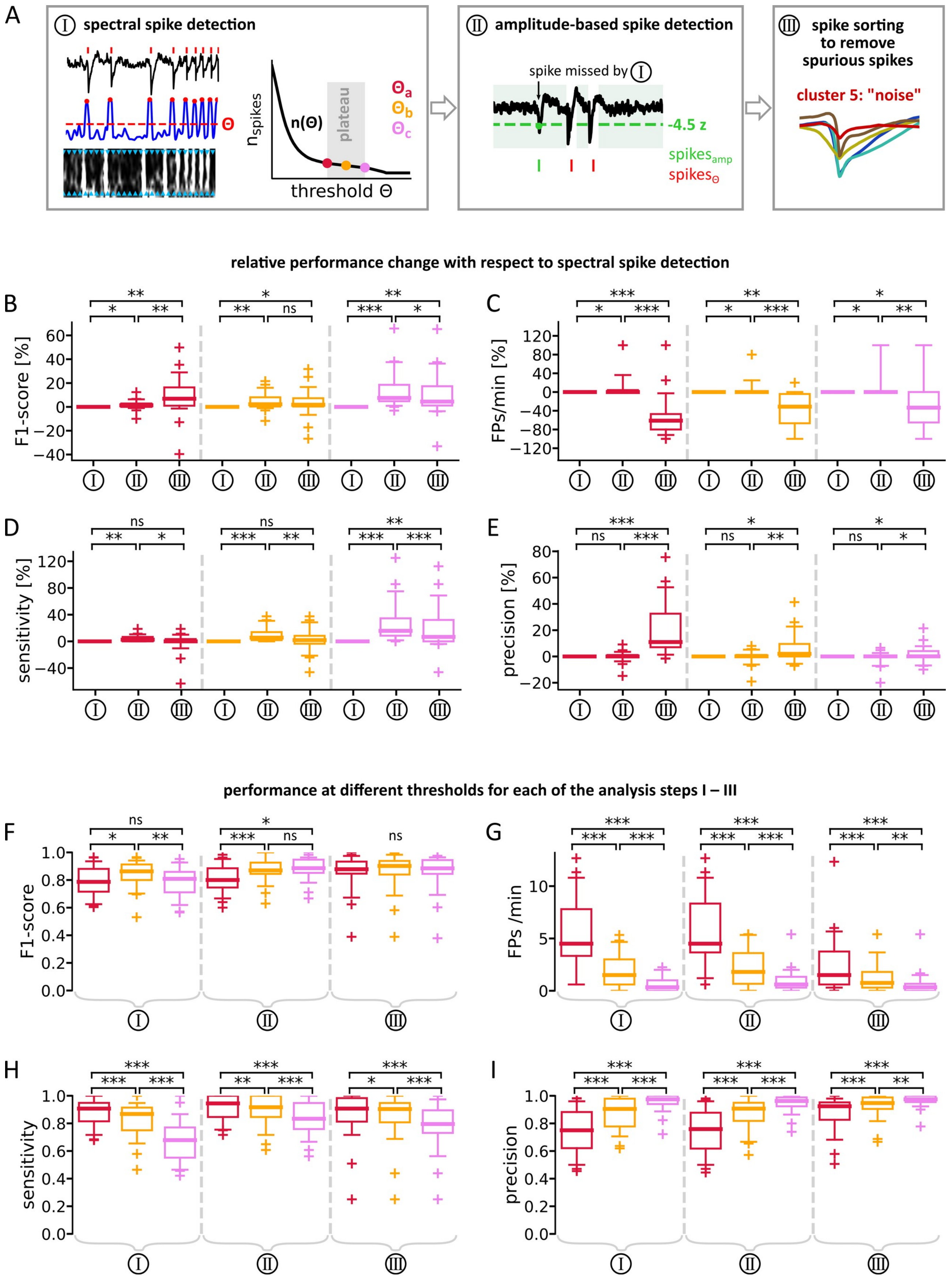


**Figure 3—figure supplement 1: Performance gains in spike detection through steps II and III**

**(A)** Summary of the spike detection procedure. Step I: Local minima of the normalized spectral sum $\widetilde{A}(t,f)$ (blue) above a certain threshold Θ (red line) are detected as $spikes_Θ$ (red dots). The candidate thresholds $Θ_a$ , $Θ_b$ and $Θ_c$ (colored dots, right) are located at the beginning, middle and end of the plateau region of n(Θ). Step II: Large amplitude spikes missed by step I are detected using an amplitude threshold (green dashed line, $spikes_{amp}$). Step III: Spike sorting is used to reduce false positive detections. Average waveforms of spike clusters are shown in colors. The spikes assigned to the cluster with the lowest peak-to-peak amplitude (red) are discarded. **(B–E)** Change in performance measures relative to step I through consecutively applying steps II and III. One sample point per recording and threshold (N = 29). Within each box, the middle bar indicates the median. Boxes enclose the $25^{th}$–$75^{th}$ percentiles. Whiskers enclose the $5^{th}$–$95^{th}$ percentiles. Crosses mark sample points outside the whisker range. Detections based on $\Theta_c$ were substantially boosted in sensitivity through step II (D), while the precision of $\Theta_a$ based detections benefited most from step III (E). Steps II and III thus balanced the respective weaknesses of $\Theta_c$ (low sensitivity) and $\Theta_a$ (low precision) based detections, while the performance of $\Theta_b$ based detections changed not as much. Applying steps II + III thus leads to overall similar F1-scores (B). Note also that step II and step III did not negatively impact false positive rate (i. e. more false positives due to amplitude detection, C) and sensitivity (i. e. spikes wrongly assigned to the noise cluster), respectively (D). **(F–H)** Comparing performance of $\Theta_a$–$\Theta_c$ separately for steps I–III. $\Theta_b$ based detections performed best when just considering step I (F). When including step II, $\Theta_c$ also became a valid choice. With adding step III, it was equally feasible to use $\Theta_a$, which excelled in terms of sensitivity (H). We conclude that $\Theta_b$ is preferable if only step I is applied. If step III, which requires the most computational resources, is omitted, we recommend using $\Theta_c$. With all three steps in combination and a focus on sensitivity, however, $\Theta_a$ should be chosen. Statistics: Kruskal-Wallis test for differences across groups followed by pairwise Wilcoxon signed-rank test with Bonferroni correction; ns $p \geq 0.05$, * $p < 0.05$, ** $p < 0.01$, *** $p < 0.001$.

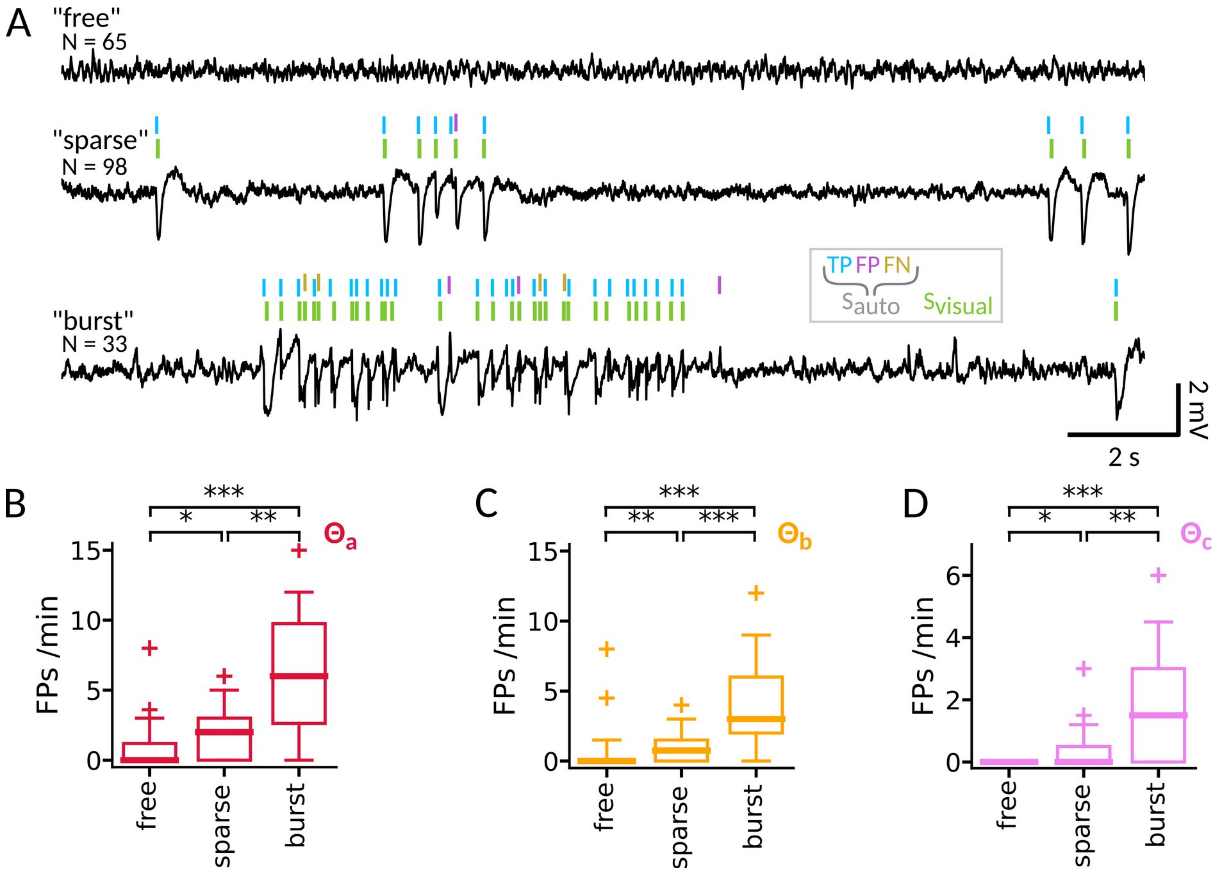


**Figure 3—figure supplement 2: Detection performance depends on context**

**(A)** Three example snippets (20 s), which an expert annotated either as spike free (*free*, top), as sparsely populated by spikes (*sparse*, middle) or as containing spikes in dense bursts (*burst*, bottom). Spikes selected by the expert are shown as green ticks ($S_{visual}$). Compared to these, spikes detected by the algorithm ($S_{auto}$) were identified as true positives (TP, blue) or false positives (FP, purple). Spikes missed by the algorithm were considered false negatives (FN, brown). In the *burst* snippet, where individual spikes are harder to distinguish, FPs and FNs are more abundant. **(B–C)** Comparison of false positive rates for detections based on $\Theta_a$ (B), $\Theta_b$ (C) and $\Theta_c$ (D). Data refers to the whole procedure (step I–III). One data point per recording session (N = 29). While false positives were comparatively abundant in *burst* snippets, they were rare in *sparse* snippets and often absent in *free* snippets. *Statistics:* Kruskal-Wallis test for differences across groups followed by pairwise Wilcoxon signed-rank test with Bonferroni correction; * $p < 0.05$, ** $p < 0.01$, *** $p < 0.001$.

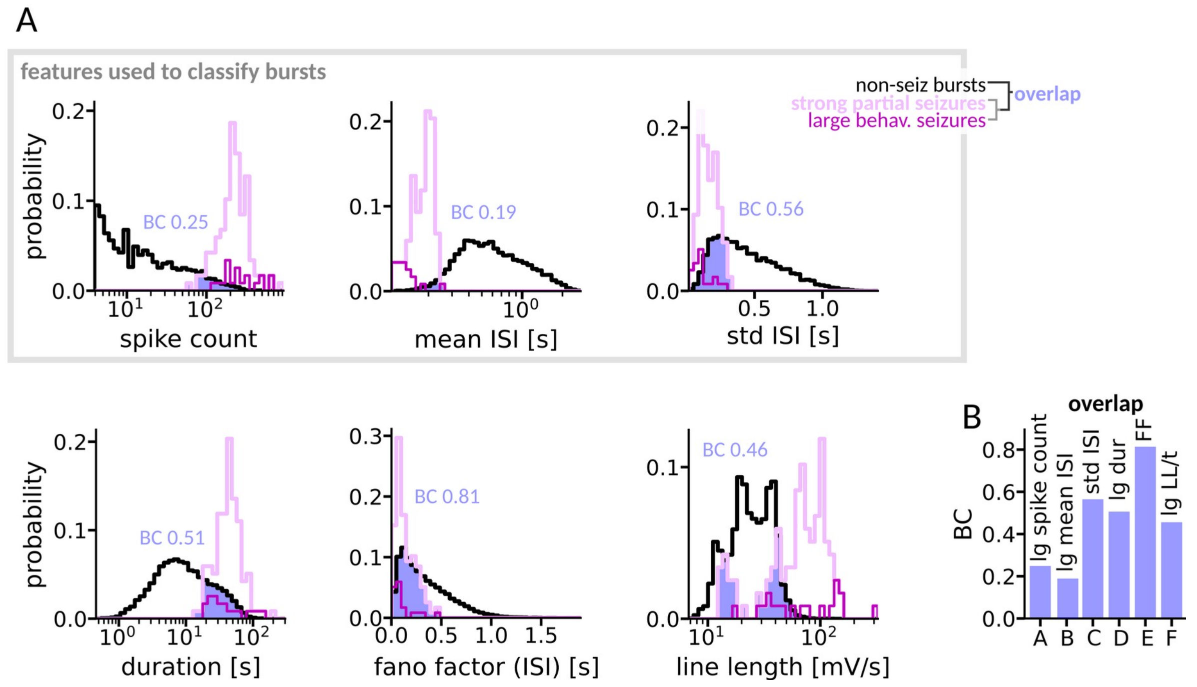


**Figure 4—figure supplement 1: Distributions of candidate features**

**(A)** Histograms of burst features. Data from bursts corresponding to visually identified seizures are shown in pink (strong partial seizures, N = 99) and magenta (large behavioral seizures, N = 18). Data from all other bursts containing at least five spikes (N = 12373) is shown in black. Histograms of bursts and seizures (behavioral and partial jointly) were each normalized to an area of 1. The overlap between seizure and non-seizure data (blue area) was quantified with the Bhattacharyya coefficient (BC, blue numbers). The gray box marks the features used for classification (also shown in Figure 4B). **(B)** Comparison of BCs given in (A). Note that *lg spike count* and *lg mean ISI* yield superior separability (low BC) compared to *lg LL/t* and *lg duration*. Abbreviations: dur = duration; FF = fano factor (ISI); LL/t = line-length/time.

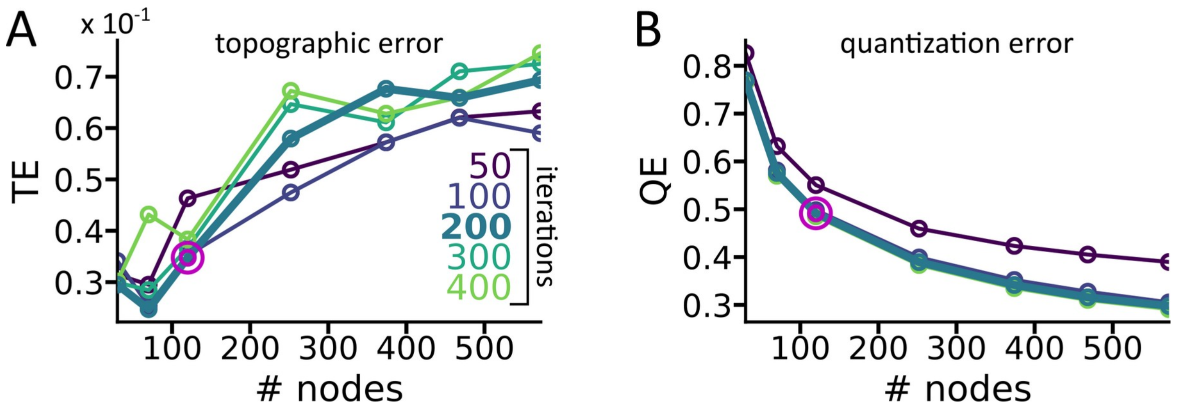


**Figure 4—figure supplement 2: SOM - Parameter choice and quality assessment**

Quality measures as a function of the number of nodes used to construct the SOM. Lines are colored according to the number of iterations with which the prototype vectors of the SOM were updated. **(A)** The topographic error gives the fraction of all input vectors for which the first and second BMU are not neighbors on the SOM. It increases with the size of the SOM. A low topographic error indicates that the topology of the input space is preserved on the SOM. **(B)** The quantization error gives the mean distance between each input vector and its BMU and thus rewards a detailed representation of the input space. Consequently, larger SOMs tend to have lower quantization errors. Chosing 120 ($6 \times 20$) nodes (magenta circles) with 200 iterations resulted in a trade-off between topology preservation and a close representation of the inputs. For identifing sub-patterns of electrographic seizures a more fine-grained classification might have been appropriate. For this study, however, we opted for a rather small SOM to keep the visual display compact and to allow faster projections of new datasets. For details about topographic and quantization errors in quality assessment see Pölzlbauer, (2004).

Pölzlbauer G. 2004. Survey and Comparison of Quality Measures for Self-Organizing Maps. *Proc Fifth Work Data Anal WDA’04* 67--82.

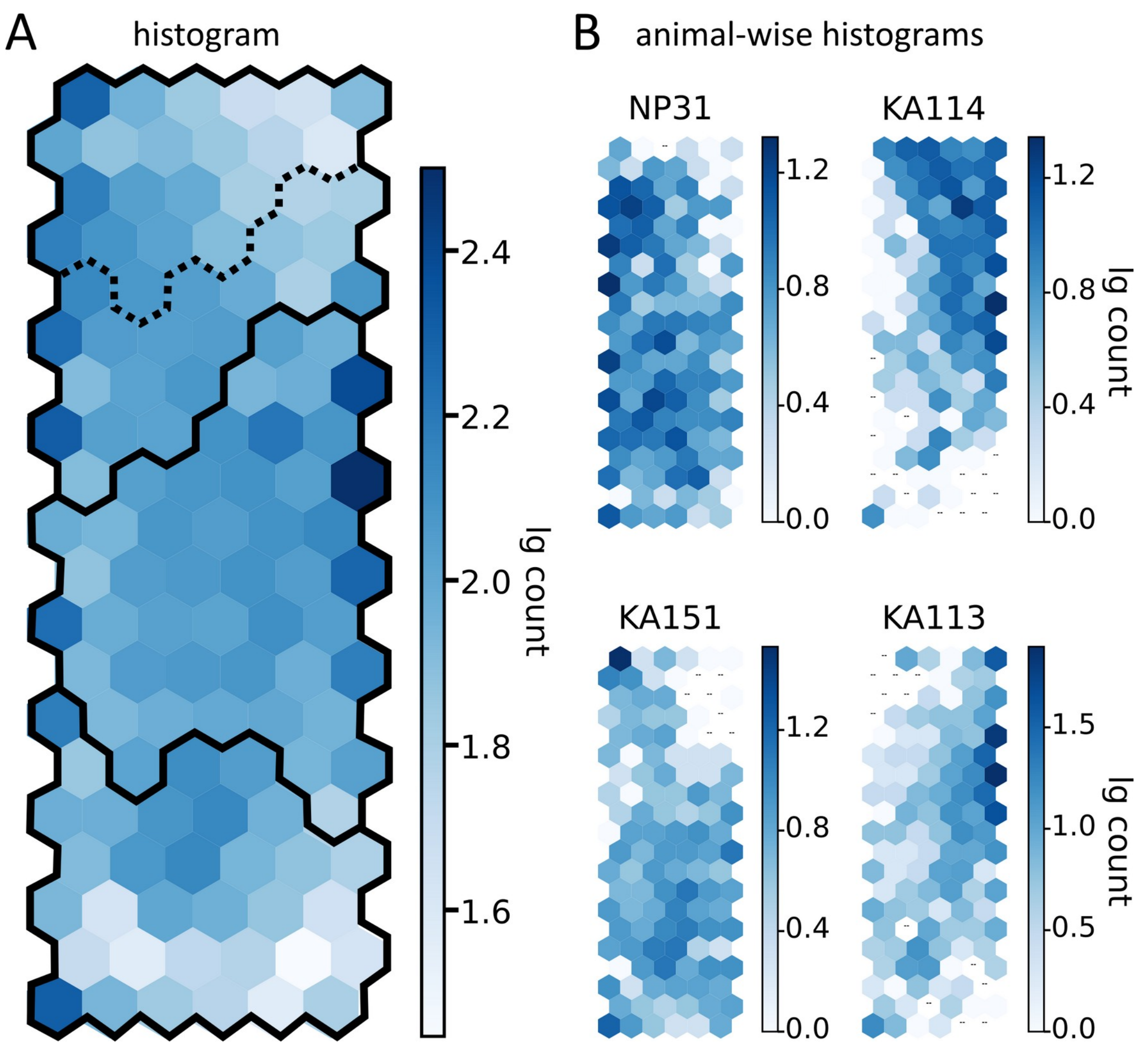


**Figure 4—figure supplement 3: Hit histogram of the SOM**

**(A)** Number of bursts matched to each node (blue colors). **(B)** Number of bursts matched to each node for bursts originating from four different example mice. Dashes indicate empty nodes. While for some mice the distributions across the map were rather even, bursts from other mice were more concentrated in certain regions of the SOM. These tendencies were in line with our visual assessment of the raw traces of the respective mouse. The sub-regions of the SOM were thus not primarily reflecting differences between mice. Instead, the occupancy of the SOM represented the differences in EA composition of different mice.

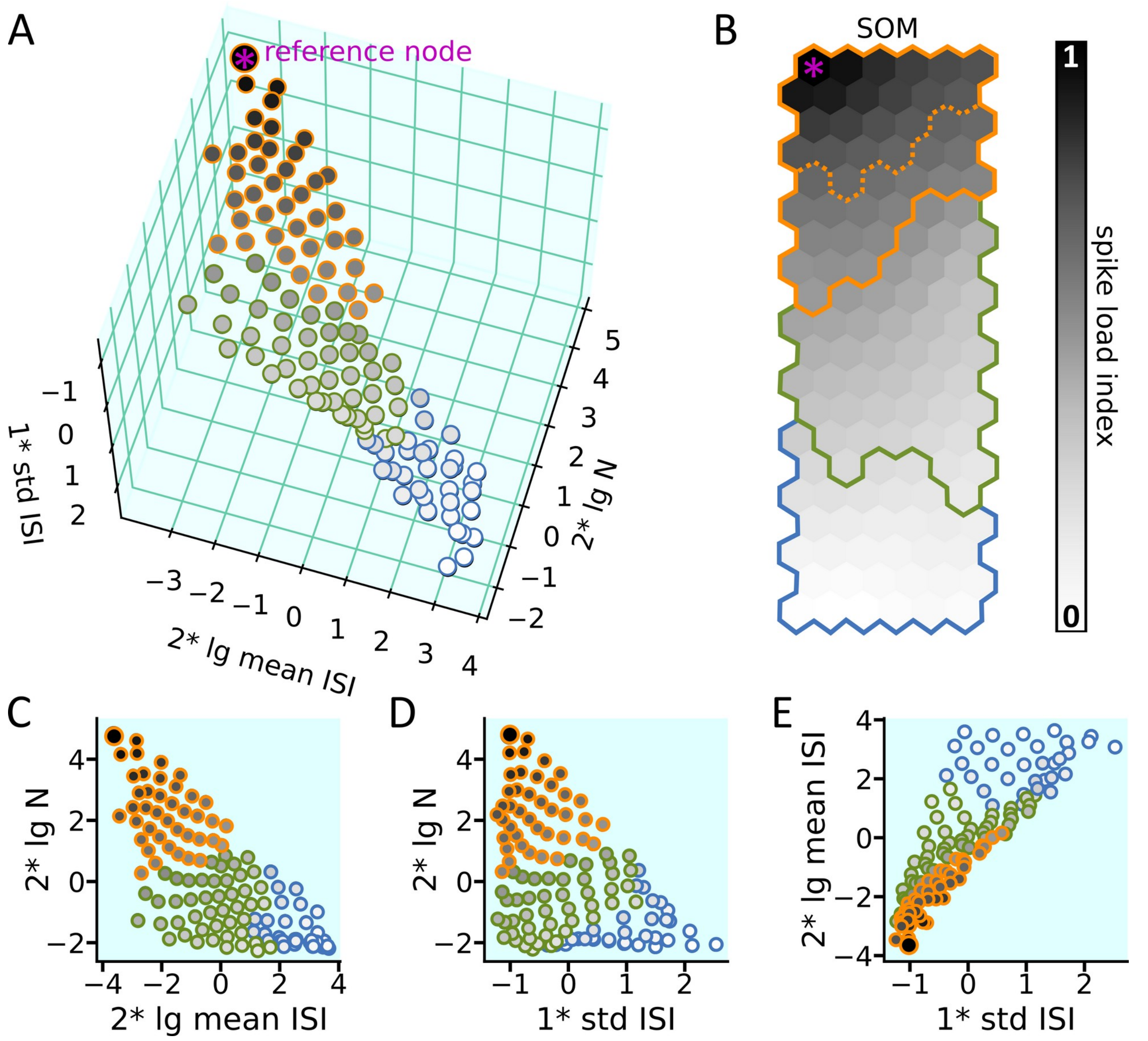


**Figure 4—figure supplement 4: The SOM's prototype vectors in feature space**

**(A)** Dots represent nodes of the SOM in feature space and are colored according to their spike load index (gray scale), which is defined by the closeness to the reference node H* (asterisk). Edge-colors indicate the categories assigned to the nodes. **(B)** SOM with nodes colored according to spike load index. Colored lines mark the borders of the categories identified in Figure 5A. **(C–E)** Two dimensional projections of (A). The spike load index displays a smooth gradient. The prototype vectors of the SOM are distributed rather continuously on a sheet in feature space. The four nodes with the highest spike load appear to be slightly set apart from the other nodes, yet no clear clusters are discernible.

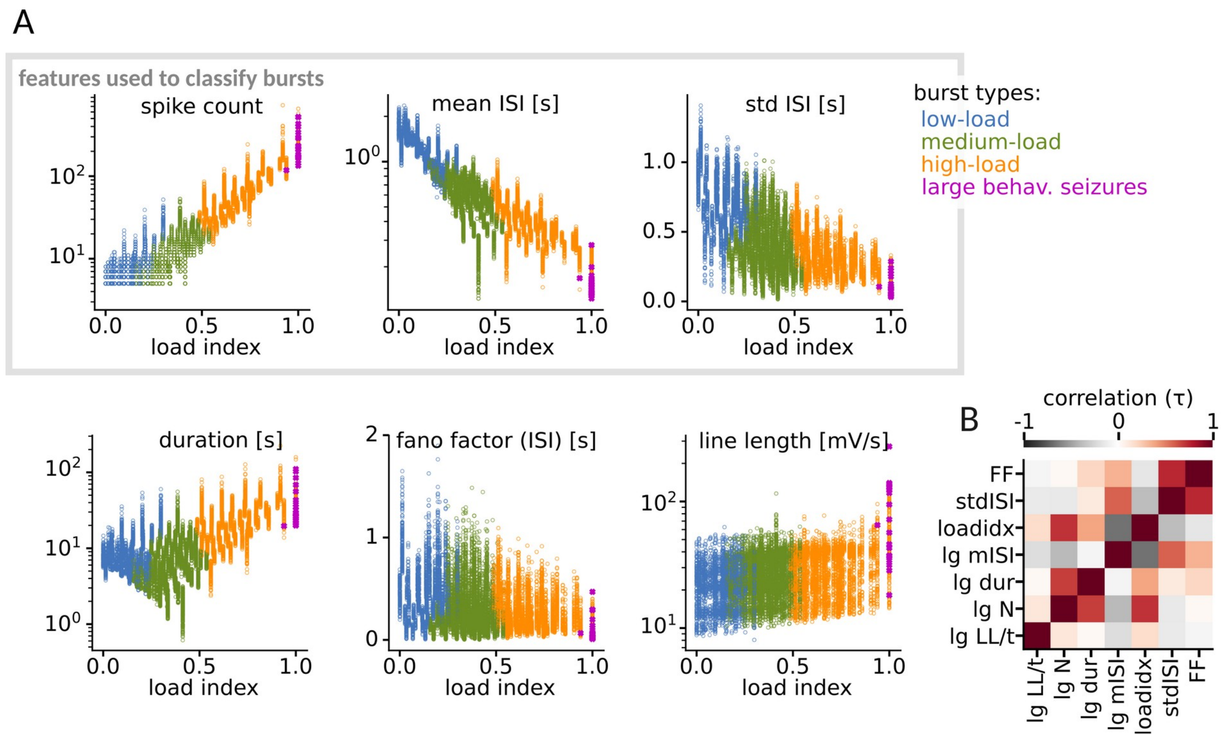


**Figure 5—figure supplement 1: Correlations between features and spike load index**

**(A)** Each panel displays the correlation between one of the candidate features shown in *Figure 4—figure supplement 1A* and the spike load index. One dot represents one burst and is colored according to its category as defined in Figure 5A. Magenta colored dots mark visually identified large behavioral seizures which were all part of the category high-load. **(D)** Correlation matrix of the features in A. All correlations were statistically significant with $p_{\tau} < 0.001$. Abbreviations: fano = fano factor (ISI); mISI = mean ISI; dur = duration; N = spike count; LL/t = line-length/time.

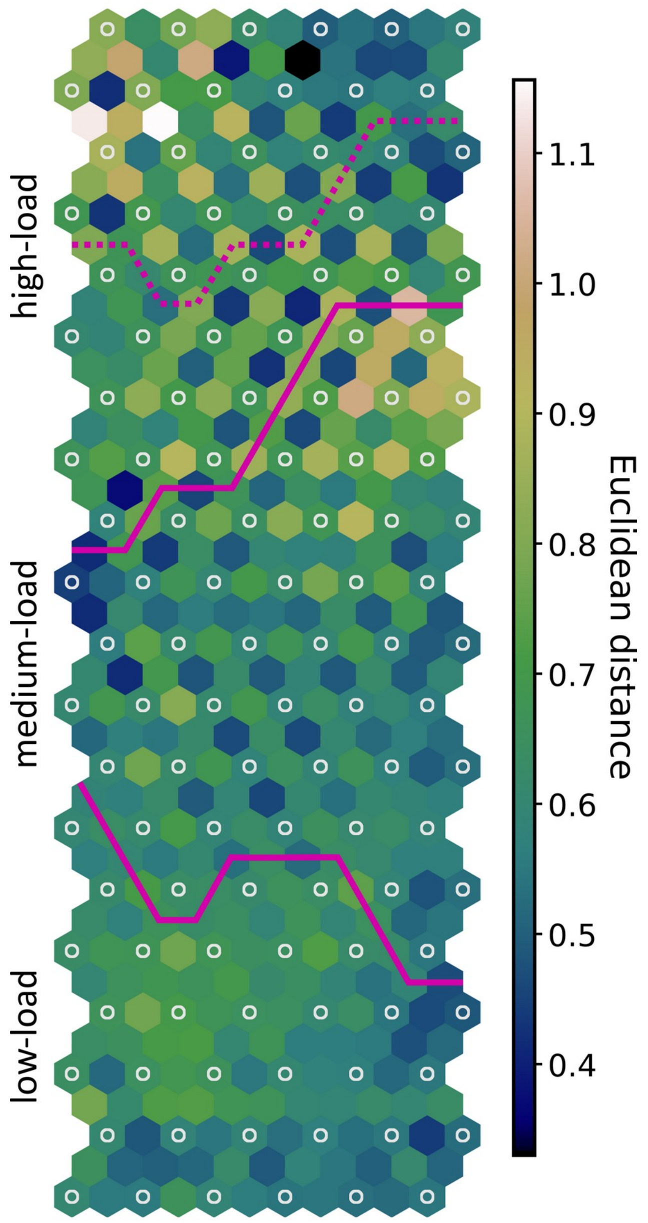


**Figure 5—figure supplement 2: The U-matrix indicates no distinct clusters in the SOM**

The U-matrix displays the Euclidean distance between prototype vectors of adjacent nodes on the SOM (Vesanto and Alhoniemi, 2000). White circles mark the original nodes. The others, so-called *interpolating nodes,* are colored according to the distance between neighboring original nodes. The color of an original node indicates the mean distance of its surrounding interpolating nodes. Borders of the categories assigned according to Figure 5 are shown as magenta lines. The top-left rows of the SOM, which represented some of the most seizure-like bursts, were clearly set apart from their diagonal neighbors located at their respective bottom-right, while they were closer to their neighbors in the horizontal and diagonal bottom-left direction. The resulting checkerboard pattern was due to the top-left to bottom-right gradient of the spike count feature that dominated the top part of the SOM. In the middle part of the SOM the pattern was reversed (bottom-right neighbors get closer) because there was a strong top-right to bottom-left gradient in the mean ISI feature

(Figure 4C). Note that the borders of the categories assigned with hierarchical clustering do not appear to match any topographical borders in the U-matrix. Still, the borders of the categories are perpendicular to the dominant gradients (top-right to bottom-left in the top part and top-left to bottom-right in the bottom part), indicating that the hierarchical clustering followed the structure of the SOM.

Vesanto J, Alhoniemi E. 2000. Clustering of the self-organizing map. *IEEE Trans Neural Networks* 11:586–600. doi:10.1109/72.846731

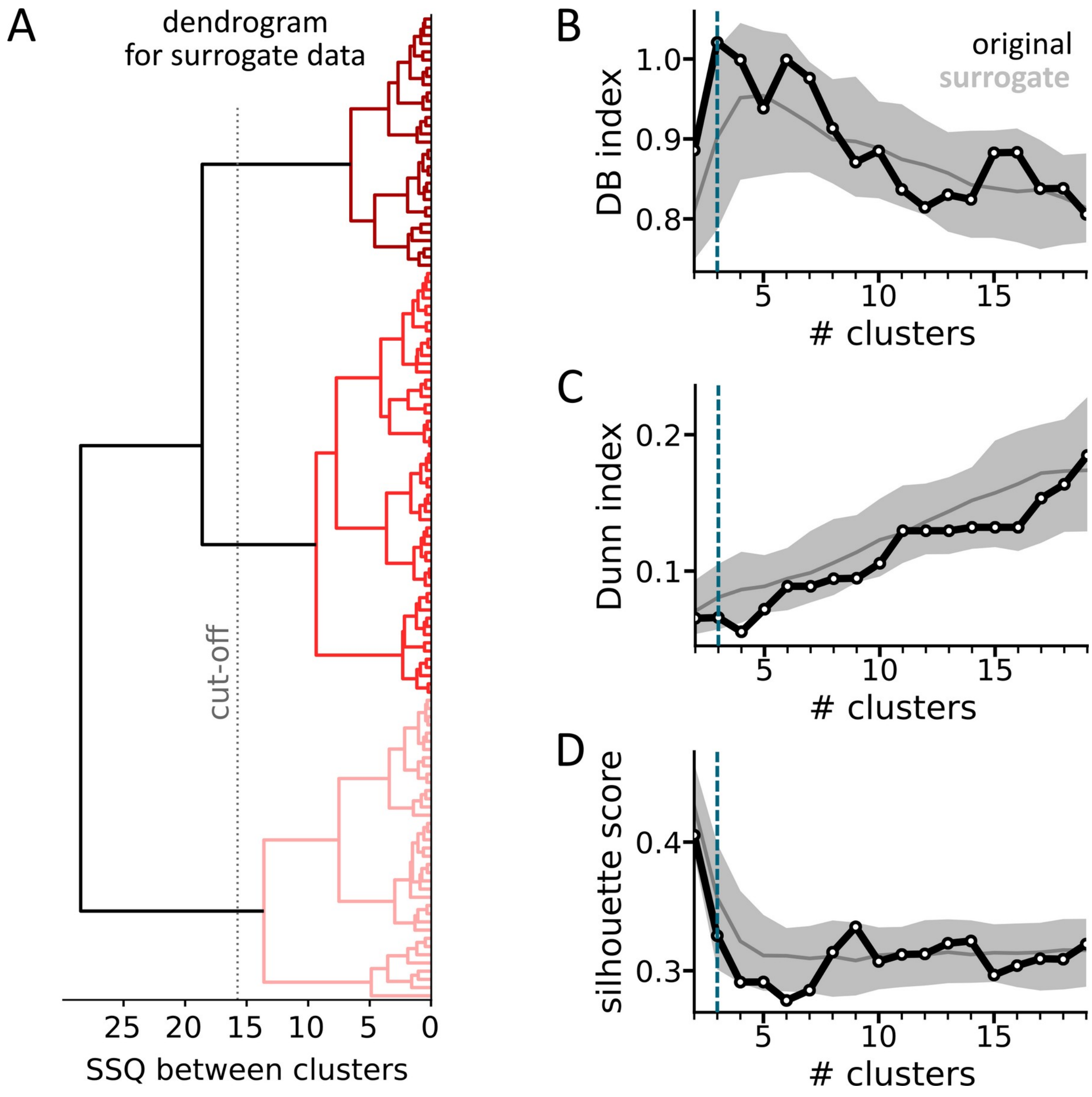


**Figure 5—figure supplement 3: The prototype vectors of the SOM are not more clustered than random samples**

**(A)** Example dendrogram generated from a surrogate set of vectors which had the same covariance structure as the original prototype vectors. With a cut-off as indicated by the gray dashed line, three main clusters (red colors) were obtained. Visually, surrogate dendrograms appeared to be very similar to the original dendrogram (compare Figure 5A). To generate a surrogate set, three independent normal distributions with 120 samples each (mean = 0, std = 1) were randomly assembled into 120 three-dimensional surrogate vectors. Using a technique based on Cholesky decomposition (Myers, 1989)), a surrogate set was then transformed to have the same covariance matrix R as the original prototype vectors. In this way, 100 surrogate sets and their dendrograms were obtained. **(B)** The black line shows the Davies-Bouldin (DB) index as a function of the number of clusters obtained from the original prototype vectors. The DB index indicates the ratio between within-cluster variability and between-cluster distance and a lower DB index indicates better clustering (Davies and Bouldin, 1979). Higher numbers of clusters were obtained by cutting the dendrogram shown in Figure 5A at decreasing levels. The gray area encloses the 10$^{th}$–90$^{th}$ percentile of DB indices obtained from the dendrograms of 100 surrogate sets, the gray line shows their median. The blue vertical line at N = 3 marks the number of clusters chosen to partition the SOM. **(C)** The Dunn index is the ratio of the smallest distance between vectors not belonging to the same cluster and the largest within-cluster distance (Dunn, 1974). A higher Dunn index indicates better clustering. **(D)** The silhouette score reflects how similar samples are on average to their assigned clusters compared to other clusters (Rousseeuw, 1987). A higher silhouette score indicates better clustering.


Davies DL, Bouldin DW. 1979. A Cluster Separation Measure. *IEEE Trans Pattern Anal Mach Intell* **PAMI-1**:224–227. doi:10.1109/TPAMI.1979.4766909

Dunn JC. 1974. Well-separated clusters and optimal fuzzy partitions. *J Cybern* **4**:95–104. doi:10.1080/01969727408546059

Myers DE. 1989. Vector conditional simulationJournal of Chemical Information and Modeling. pp. 283–293. doi:10.1007/978-94-015-6844-9_21

Rousseeuw PJ. 1987. Silhouettes: a graphical aid to the interpretation and validation of cluster analysis. *J Comput Appl Math* **20**:53–65. doi:10.1016/0377-0427(87)90125-7

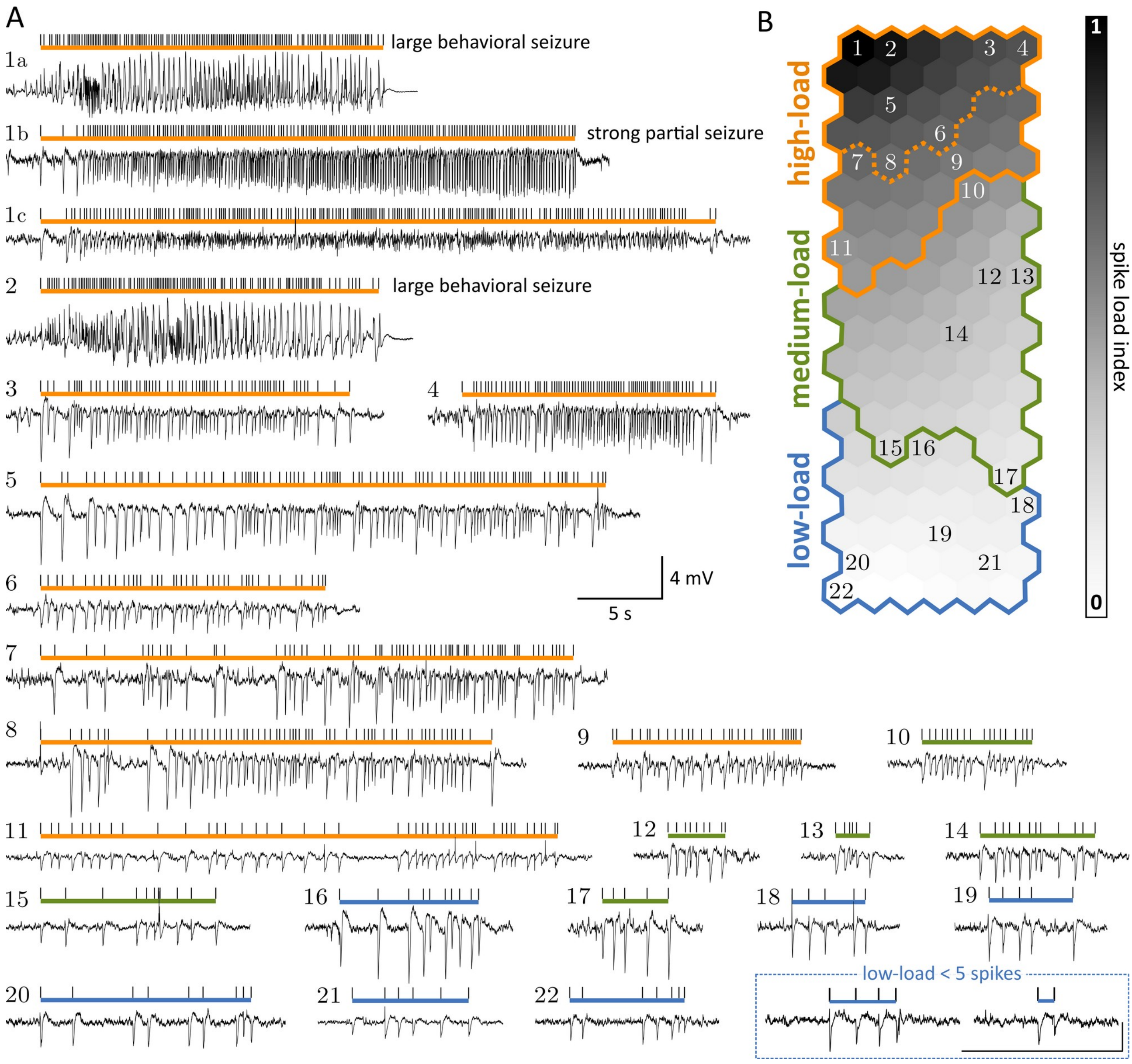


**Figure 5—figure supplement 4: LFP morphology of bursts assigned to different nodes on the SOM**

**(A)** Exemplary bursts. The colored bars indicate the extent and category of a respective burst, black ticks mark spikes. The numbers refer to the location of a burst's BMU on the SOM shown in B. The two bursts in the bottom right have fewer than five spikes, were not classified on the SOM, and were categorized as low-load. **(B)** SOM with nodes colored according to spike load index. Colored lines indicate the borders of categories. Numbers refer to examples shown in A. Both panels were adapted from Heining et al. (2019).
